# Epizootic tipping points: Environmental viral feedbacks predict amphibian die-offs

**DOI:** 10.64898/2026.03.24.714032

**Authors:** Logan S. Billet, Jason T. Hoverman, Erin L. Sauer, John-Gabriel Bermudez, David K. Skelly

## Abstract

Virulent pathogens often circulate in wildlife populations without causing mass mortality, but what distinguishes outbreaks that escalate to die-offs remains poorly understood. We tracked ranavirus epizootics across 40 wood frog breeding ponds over three years, where infection was widespread but die-offs occurred in only 13 of 102 pond-years. Models based on viral state received greater support than abiotic and host conditions for subsequent infection prevalence, viral load, and environmental viral concentration (eDNA). Infection prevalence predicted later eDNA concentration throughout epizootics, consistent with continued shedding from infected hosts. In contrast, relationships between eDNA concentration and subsequent infection prevalence and viral load were strongly attenuated when die-off-onset visits were excluded. Dynamic viral trajectories distinguished die-off from non-die-off pond-years more clearly than static pond and host features. Our results point toward a phase-dependent feedback in which pathogen accumulation in the shared environment contributes to the escalation from widespread infection to mass mortality.

## Introduction

Mass mortality events can have substantial ecological, evolutionary, and economic consequences, and appear to be increasing globally (Campbell *et al*. 2018; Earl & Gray 2014; Fey *et al*. 2015, 2019; Potts *et al*. 2010). Across animal taxa, disease is a prominent driver of mass mortality (Fey *et al*. 2015), yet a central question remains largely unresolved: what distinguishes outbreaks that escalate to mass mortality from those that do not? This challenge is apparent across systems. For example, *Batrachochytrium dendrobatidis* can persist enzootically at low infection loads yet drives rapid population collapse when infection intensity exceeds a critical threshold (Briggs *et al*. 2010; Vredenburg *et al*. 2010), and white-nose syndrome causes mass mortality in some bat hibernacula while colonies of the same species persist at others (Frick *et al*. 2010; Langwig *et al*. 2012). In these and other systems, infection is often widespread, but what determines whether widespread infection leads to mass mortality remains poorly understood.

Predominant approaches to explaining mass mortality have focused on individual- level susceptibility and identifying the environmental and physiological conditions that make individual hosts more likely to become infected, develop severe disease, and/or die from infection (Beldomenico & Begon 2010; Brearley *et al*. 2013; Hing *et al*. 2016; Rollins-Smith 2017). These approaches have been productive across wildlife disease systems, yielding a strong understanding of how temperature, nutritional stress, coinfection, and immune status shape individual infection outcomes. Ranaviruses, among the most significant pathogenic threats to amphibians globally (Gray *et al*. 2009; Price *et al*. 2014; Teacher *et al*. 2010), have received considerable attention in this context. Lab studies have identified many factors that impact individual infection outcomes, including temperature (Brand *et al*. 2016; Echaubard *et al*. 2014), developmental stage (Haislip *et al*. 2011), and environmental stressors (Forson & Storfer 2006; Hall *et al*. 2020), and field studies have suggested that the seasonal timing of die-offs is consistent with temperature- and stage-dependent shifts in pathogenicity (Hall *et al*. 2018). The implication for population-level outcomes is that if individual susceptibility depends on identifiable environmental conditions, those same conditions should best predict when and where populations are at risk of severe disease outbreaks (Lafferty & Holt 2003).

However, population-level outcomes have not cleanly followed these individual- level predictions (Brunner *et al*. 2025b). In controlled experiments, conditions commonly considered stressful, including predation risk, crowding, food limitation, and chemical contaminants, have not substantially increased the severity of ranavirus epizootics (Haislip *et al*. 2012; Jacobson 2019; Reeve *et al*. 2013). This pattern extends to the stressors with some of the strongest documented effects on infection outcomes; well-replicated manipulations of temperature and salinity, which strongly influence the intensity and lethality of ranavirus infections in individual tadpoles (Brand *et al*. 2016; Hall *et al*. 2020), had minimal influence on the probability, severity, or timing of epizootics in mesocosm populations (Brunner *et al*. 2025a). These results suggest that the transition from widespread infection to mass mortality depends on processes not fully captured by variation in individual host susceptibility. Thus, individual susceptibility may shape which hosts become infected and how severely, but population-level processes may help determine whether infection escalates to mass mortality.

One such process is the accumulation of pathogens in the shared environment, which shifts the exposure landscape experienced by the population rather than host susceptibility. Die-off events in wildlife are disproportionately concentrated in aquatic systems (>90% reported in Fey *et al*. 2015), where water exchange may be limited, pathogen dispersal from shedding hosts may be spatially constrained, and waterborne pathogens can persist outside hosts for extended periods (Murray 2009; Nazir *et al*. 2012). In such systems, infected hosts shed pathogens into a shared medium, and when shedding outpaces decay, pathogen concentrations can rise over time. For many host-pathogen systems, including ranavirus, pathogen dose influences both whether infection establishes and how severe infection becomes (Brunner *et al*. 2005; Forzán *et al*. 2017; Lunn *et al*. 2019; Warne *et al*. 2011), and the dose an individual experiences is often not fixed at the point of initial exposure. When infected hosts shed into a shared reservoir, the collective output of the infected population shapes the ongoing exposure dose each host experiences, creating the potential for a population-level positive feedback: rising pathogen concentrations intensify infection, increasing shedding and further elevating pathogen concentrations.

Many existing models of environmentally transmitted pathogens connect shared reservoirs primarily to the force of infection but not to disease progression within already-infected hosts (Briggs *et al*. 2010; Codeço 2001; Tien & Earn 2010). The feedback described above operates through both. Virus in the environment drives new infections and intensifies existing ones, generating dynamics that these frameworks do not fully capture. The conditions that may permit such a feedback, including environmental pathogen persistence, dose-dependent disease severity, and limited acquired immunity, are documented in the ranavirus system (Andino *et al*. 2012; Brunner & Yarber 2018; Nazir *et al*. 2012; Rollins-Smith 1998), where accumulation rate should depend on host density, shedding rates, water volume, and viral decay. Under this framework, the key question shifts from what makes individual hosts susceptible to what governs pathogen accumulation in the shared environment. We formalize this mechanism in a compartmental model (Billet *et al*. 2026; Fig. 1); here, we test whether the feedback structure it describes is detectable in natural populations.

**Figure 1.**
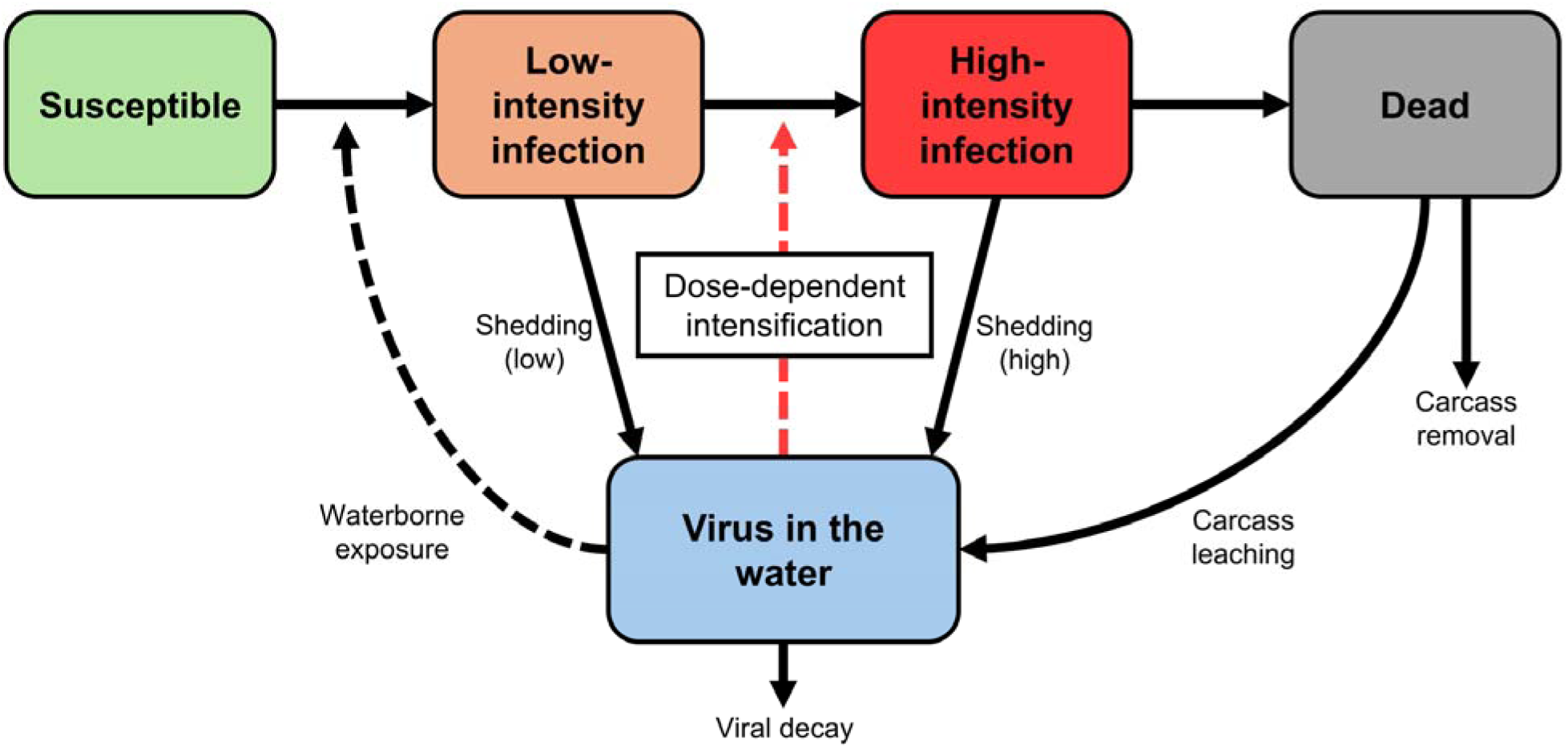
**Hypothesized environmental viral feedback underlying ranavirus die-offs**. Susceptible hosts acquire infection through exposure to ranavirus in the shared water column, initially entering a low-intensity infection state. Increasing environmental viral exposure promotes both new infections and dose-dependent progression from low-to high-intensity infection. Infected hosts shed virus into the water, with greater shedding from high-intensity infected hosts and carcasses providing an additional viral input. Environmental viral accumulation represents the balance between these inputs and losses via viral decay and carcass removal. Environmental DNA concentration provides a field measure of viral genetic material in the shared water column.

We tracked ranavirus epizootics across 40 wood frog (*Rana sylvatica*) breeding ponds over three years, measuring individual infection status and viral load alongside environmental viral concentrations (eDNA) at biweekly intervals. We first asked what predicted tissue-confirmed infection occurrence at the pond-year level. Then, for three within season responses— subsequent visit-level infection prevalence, viral load among infected individuals, and environmental viral concentration—we compared models based on lagged viral state variables against models based on abiotic and host predictors. If pathogen accumulation in the shared environment governs epizootic outcomes, we predict (1) that lagged viral state variables will be stronger predictors of these within-season responses than environmental and host conditions. If the positive feedback described above operates, we predict (2) that lagged eDNA concentration and infection prevalence should each predict the other’s subsequent state: higher infection prevalence should predict higher subsequent waterborne viral concentration, and higher environmental viral concentration should predict higher subsequent infection prevalence and viral loads. Finally, if this feedback contributes to the transition from widespread infection to mass mortality, we predict (3) that the relationship between environmental virus and subsequent host infection should be strongest near die-off onset. Specifically, environmental viral concentration should predict higher subsequent infection prevalence and viral loads when die-off onset visits are included, but these relationships should be attenuated and lose statistical support when onset visits are excluded. Our results support these predictions, suggesting that epizootic outcomes may be shaped by a self-reinforcing dynamic operating through the shared viral environment.

## Material and methods

### Study system and field sampling

From 2021 through 2023, we surveyed wood frog breeding ponds in Yale-Myers Forest (YMF), a 3,213-ha forest in northeastern Connecticut, USA (Billet *et al*. 2024; Rowland *et al*. 2022) (Fig. 2A). These fishless, temporary ponds support low amphibian diversity; wood frog tadpoles comprise the vast majority of vertebrate biomass from April through July. We surveyed 30-40 ponds annually (102 pond-years), including sites with documented ranavirus die-offs (Hall *et al*. 2018, 2020).

**Figure 2.**
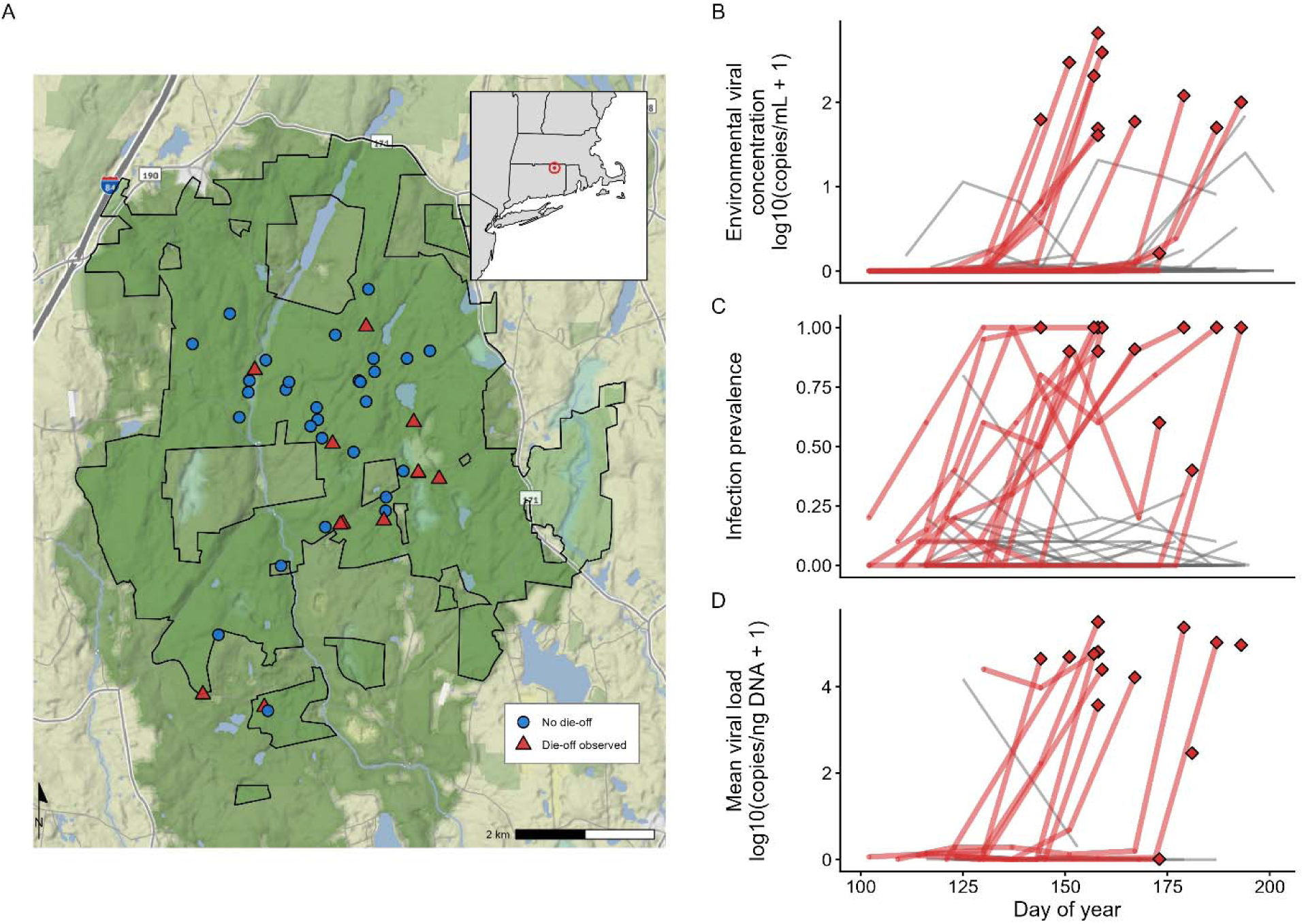
Study system and epizootic variation at Yale-Myers Forest. (A) Locations of 40 study ponds at Yale-Myers Forest, Connecticut, USA. Triangles indicate ponds in which at least one die-off was observed (n=11); circles indicate ponds with no observed die-offs. The inset shows the study location in the northeastern United States. (B-D) Pond-year trajectories across the larval season for die-off (red) and non-die-off (gray) pond-years: (B) environmental viral concentration measured using eDNA, (C) infection prevalence, and (D) mean viral load among infected individuals. Diamonds indicate the visit at which a die-off was first detected. Values in B and D are log_10_(x+1) transformed.

We censused egg masses within approximately one week of breeding and sampled ponds approximately every 2 weeks throughout larval development (April-July). At each visit, we estimated tadpole density via timed dip-net surveys (Werner *et al*. 2007), and haphazardly collected tadpoles with a target of 20 individuals per visit. We recorded water chemistry (conductivity, pH, dissolved oxygen) and maximum depth and collected ten 100-mL surface-water subsamples around the pond perimeter, which were pooled, stored on ice, and filtered within 24 h for ranavirus eDNA quantification. Pond temperatures were logged continuously.

Static pond attributes included canopy openness, surface area, elevation, and surrounding land cover. We monitored for mortality weekly via visual encounter surveys, as ranavirus die-offs in this system are characterized by rapid mortality over days to ∼2 weeks (Gray *et al*. 2009). We defined die-off onset as the first visit at which ≥5 carcasses were observed (Hall *et al*. 2018). See Appendix S1 for full study system and field sampling details.

### Laboratory methods

Collected tadpoles were staged (Gosner 1960), measured, and liver tissue dissected for DNA extraction (Qiagen DNeasy kits; Qiagen, Hilden, Germany). We screened 2,685 of 7,955 collected tadpoles via primer-probe quantitative PCR (qPCR) targeting the ranavirus major capsid protein (Leung *et al*. 2017) to assess infection status and estimate viral load (viral copies/ng DNA). We classified tadpoles as infected when amplification was confirmed in at least two qPCR reactions (Appendix S1). Because processing all tadpoles was not feasible, individuals submitted for qPCR were selected randomly from the collected pool within each focal visit. We targeted at least 30 individuals across three visits per pond-year and increased screening effort during die-offs to resolve rapid changes in prevalence and viral load. Ranavirus eDNA concentration was quantified from filtered water samples using the same primer-probe set (viral copies/mL). Laboratory deionized-water process blanks carried through filtration, extraction, and qPCR, together with extraction and plate negatives, never amplified. See Appendix S1 for full laboratory methods.

### Data structure and variable definitions

Our data comprised individuals nested within visits and pond-years. Our primary response variables were tissue-confirmed ranavirus occurrence (≥1 tissue-positive tadpole detected in a pond-year), visit-level infection prevalence (the proportion of screened tadpoles that were tissue-positive), viral load among infected tadpoles (viral copies/ng DNA), and environmental viral concentration measured by eDNA (copies/mL). We analyzed visit-level eDNA detection separately from environmental viral concentration among eDNA-positive visits. Because observed viral loads were strongly bimodal, we also analyzed viral load as a binary response, with each infected tadpole classified as a low-or high-intensity infection based on a two-component Gaussian mixture model (Appendix S1).

For within-season analyses, responses at visit *t* were related to viral conditions measured at the preceding visit (∼2 weeks earlier): eDNA concentration, infection prevalence, and, where relevant, mean viral load. For first visits, lagged prevalence was set to zero and lagged eDNA to ln(0 + 0.0001). Visits following unscreened visits were excluded from analyses requiring lagged prevalence or viral load rather than bridging across sampling gaps. Primary models used concurrent visit-level temperature, conductivity, pH, and depth, with an additional analysis giving these predictors the same one-visit lag. Temperature was summarized as a 7-day mean; residuals of temperature and developmental-stage represented deviations from their seasonal relationships with day of year. Additional predictors included day of year, tadpole density, egg mass counts, and static pond attributes (Appendix S1).

### Analytical framework

We organized analyses around four questions: (1) what predicted tissue-confirmed ranavirus occurrence; (2) whether recent viral state or environmental and host conditions better predicted subsequent infection prevalence, viral load, and environmental viral concentration; (3) whether the six directional time-lagged relationships among eDNA concentration, prevalence, and mean viral load supported the predicted feedback when evaluated individually and simultaneously using piecewise SEM; and (4) whether viral trajectories distinguished die-off from non-die-off pond-years and the same viral-state relationships appeared in an independently collected dataset from the same forest. Inference rested on convergence across these analyses rather than any single test.

We used AICc (Burnham & Anderson 2002) to compare candidate predictor classes fit to common analytical datasets, marginal R^2^ (Nakagawa & Schielzeth 2013) for fixed-effect explanatory power, and leave-one-pond-year-out cross-validation for out-of-sample predictive accuracy. Mixed-effects models used random intercepts appropriate to each sampling level (Table S1.2), and prevalence models included observation-level random effects for extra-binomial variation (Harrison 2014).

Primary within-season models included die-off-onset visits but excluded post-onset observations. We then excluded onset visits to test whether these relationships persisted during pre-die-off buildup or weakened when onset observations were removed. Our interpretation focused on changes in effect estimates and uncertainty. Additional sensitivities tested lagged environmental predictors, raw versus residualized temperature, and reduced per-visit screening.

Each directional time-lagged test asked whether one state variable at *t*−1 predicted another at *t* after accounting for the response’s prior value and day of year. We tested all six directional pairings among eDNA concentration, prevalence, and mean viral load, with P-values Holm-adjusted across the six tests. We then used piecewise SEM (Lefcheck 2016) to evaluate these pathways simultaneously, retaining the most parsimonious structure with adequate fit, and repeated the analysis without die-off onset visits (Appendix S1).

### Statistical analysis

All analyses were conducted in R version 4.5.3 (R Core Team 2026). We report 95% confidence intervals for focal regression coefficients, with computation methods described in Appendix S1. We modeled tissue-confirmed ranavirus occurrence at the pond-year level using binomial GLMMs with pond random intercepts (lme4; Bates *et al*. 2015). Candidate predictors were compared on a common complete-case sample, and the selected model was then refit using all pond-years with data for its retained predictors. We tested spatial structure in ranavirus occurrence, die-off occurrence, and occurrence-model residuals using Moran’s I (spdep; Bivand *et al*. 2026) , and tested distance to the nearest die-off pond as an additional spatial predictor.

Within-season infection prevalence, viral load, and environmental viral concentration were analyzed using the model-comparison framework described above. We analyzed visit-level eDNA detection separately (Appendix S1).

For die-off discrimination, we used Firth-penalized logistic regression (Heinze & Schemper 2002), appropriate for the small number of events (13 die-offs across 102 pond-years). Candidate predictors were static pond features and dynamic viral trajectory metrics, including pond-year-level slopes, peaks, and cumulative eDNA. Metrics were computed from pre-onset visits for die-off pond-years and from the full season for non-die-off pond-years. Because these windows differed between groups, we primarily interpret trajectory analyses as retrospective discrimination rather than prospective prediction. To ask whether these trajectories were themselves predictable from measured conditions, we also modeled prevalence and eDNA slopes as continuous responses against static environmental, first-detection, and seasonal predictors (Table S2.10). We evaluated die-off discrimination using receiver operating characteristic curves (pROC; Robin *et al*. 2011) with cluster-bootstrapped 95% confidence intervals and assessed the stability of Youden’s J optimal cutpoint (Appendix S1.4).

## Results

### Summary

After excluding post-die-off observations (274 individuals, nearly all infected), the analytical dataset comprised 2,411 screened individuals, of which 337 were ranavirus-positive. We quantified ranavirus eDNA from water samples at 594 visits across 102 pond-years.

Ranavirus was widespread across the forest. Viral eDNA was detected in 75-87% of ponds per year, while ranavirus infection was confirmed in at least one tadpole in 32 of 102 pond-years: 11 of 40 ponds in 2021, 12 of 30 in 2022, and 9 of 32 in 2023. This difference likely reflects eDNA’s greater sensitivity for pond-level ranavirus detection. Water samples integrate viral material shed by any infected host using the pond, including species not screened, and may detect virus when infection prevalence among tadpoles is too low to be detected among sampled individuals. We observed 13 die-off events across the study period: 8 of 40 ponds in 2021 (20.0%), 1 of 30 in 2022 (3.3%), and 4 of 32 in 2023 (12.5%), with onset dates ranging from late May to mid-July (DOY 144-193). Among infected individuals, viral loads spanned nine orders of magnitude; in non-die-off ponds and pre-die-off visits, viral load remained low (median=0.012 copies/ng DNA), whereas at die-off onset, median viral load exceeded 15,000 copies/ng DNA (Fig. 2D).

### Ranavirus occurrence

Ranavirus was detected in tissue samples from 32 of 102 pond-years. Higher water conductivity (β=0.90 [95% CI: 0.10-1.71], p=0.028) and greater egg mass counts (β=0.63 [95% CI: 0.10-1.16], p=0.020) were both associated with increased probability of tissue-confirmed ranavirus occurrence in the combined model (R^2^=0.26; Fig. S2.3). Spatial clustering in ranavirus occurrence was absent (Moran’s I=0.014, p=0.233).

### Infection prevalence

Among pond-years where infection was detected on at least one visit (’ever-infected’ pond-years; N=95 visits, 25 pond-years), viral state variables were more strongly supported than environmental predictors of visit-level infection prevalence. Lagged eDNA concentration and lagged prevalence were the only predictors retained in the best-supported model (eDNA*_t-1_* β=1.58 [95% CI: 0.76-2.40], p<0.001; prevalence*_t-1_* β=1.62 [95% CI: 0.68-2.56], p<0.001; R^2^=0.50; Fig. 3A,B; Table S2.2). Of the environmental and host predictors added individually, only developmental stage residuals improved model support by >2 AICc units (lower prevalence among tadpoles more advanced than expected for date; ΔAICc=−4.6, β=−0.91, p=0.009; Table S2.5). Water temperature was also negatively associated with prevalence but provided limited improvement in model support (warmer conditions associated with lower prevalence; ΔAICc=−0.5, β=−0.77, p=0.044). A parameterization separating eDNA detection from quantity showed that viral concentration conditional on detection, but not detection of virus alone, predicted subsequent prevalence (Fig. S2.4). When tadpole density and infection prevalence were both measured at the preceding visit, density did not modify the relationship between prior and subsequent infection prevalence (density × prevalence: β=0.11 [95% CI: −0.49–0.71], p=0.721).

**Figure 3.**
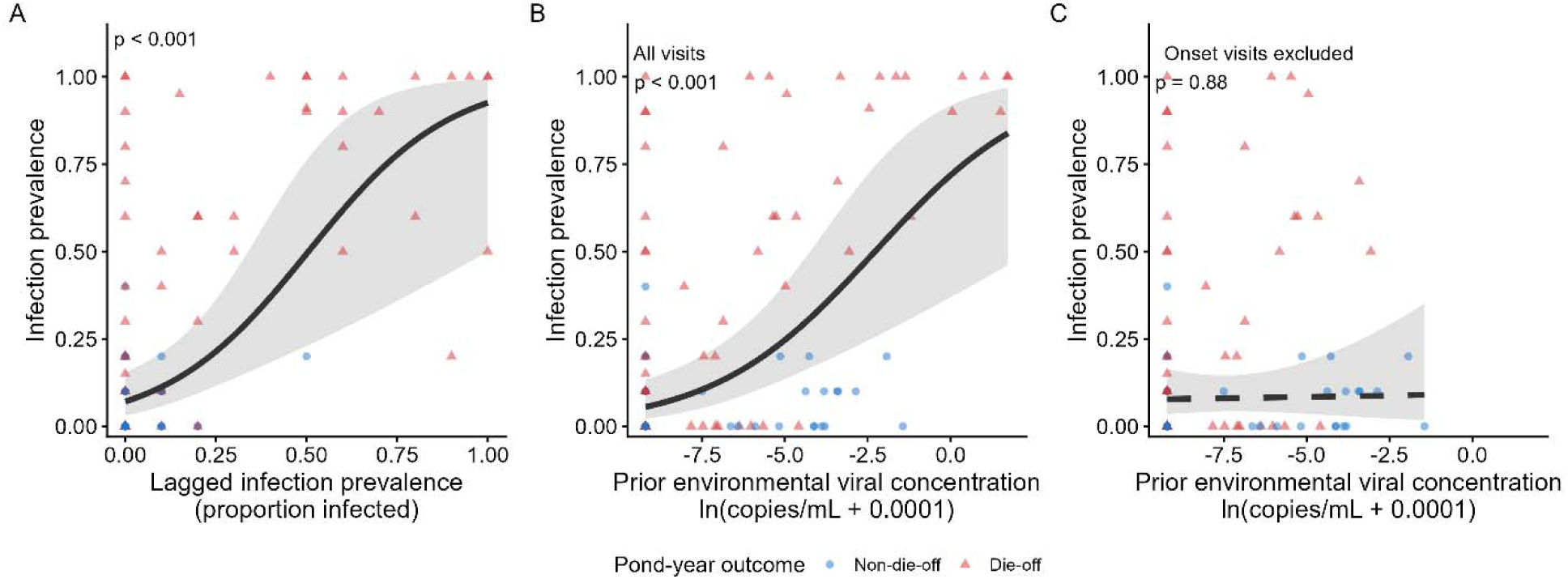
Drivers of ranavirus infection prevalence. (A) Relationship between infection prevalence at visit *t*−1 and prevalence at visit *t*, holding log-transformed environmental viral concentration at its mean among the modeled visits. (B) Relationship between environmental viral concentration at visit *t*−1 and infection prevalence at visit *t*, holding lagged prevalence at its mean among the modeled visits. (C) The relationship shown in B after excluding die-off-onset visits. Panels A and B use 95 visits; C uses 82 visits, with the non-focal mean calculated separately for that sample. Red triangles indicate die-off pond-years; blue circles indicate non-die-off pond-years. Lines show fixed-effect predictions from binomial GLMMs with pond-year and observation-level random intercepts, with random effects set to zero; shaded bands show 95% confidence intervals.

The best model retained moderate out-of-sample predictive accuracy in leave-one-pond-year-out cross-validation (LOOCV R^2^=0.40; Fig. S2.5). When tissue screening was capped at 10 randomly selected individuals per visit and iterated 500 times, both lagged eDNA and lagged prevalence remained significant in every iteration, indicating that the prevalence results were robust to unequal screening effort. Excluding die-off onset visits (N=82 visits, 25 pond-years) showed that the lagged-prevalence effect remained similar (β=1.85 [95% CI: 0.74-2.97], p=0.001), whereas the lagged-eDNA effect was strongly attenuated (β=0.08, p=0.881; Fig. 3C). Thus, prior prevalence remained associated with subsequent prevalence during pre-die-off buildup, whereas the relationship between environmental viral concentration and subsequent prevalence weakened substantially when die-off-onset visits were excluded.

### Viral load

Individual viral loads were analyzed among infected tadpoles only (i.e., uninfected individuals were excluded and not assigned zero values for viral load). In the infected subset (N=337), viral loads were strongly bimodal (Hartigan’s dip test: D=0.054, p<0.001), with low- and high-intensity components centered at approximately 0.07 and 30,000 copies/ng DNA, respectively (72.4% and 27.6% of individuals; Fig. 4A).

**Figure 4.**
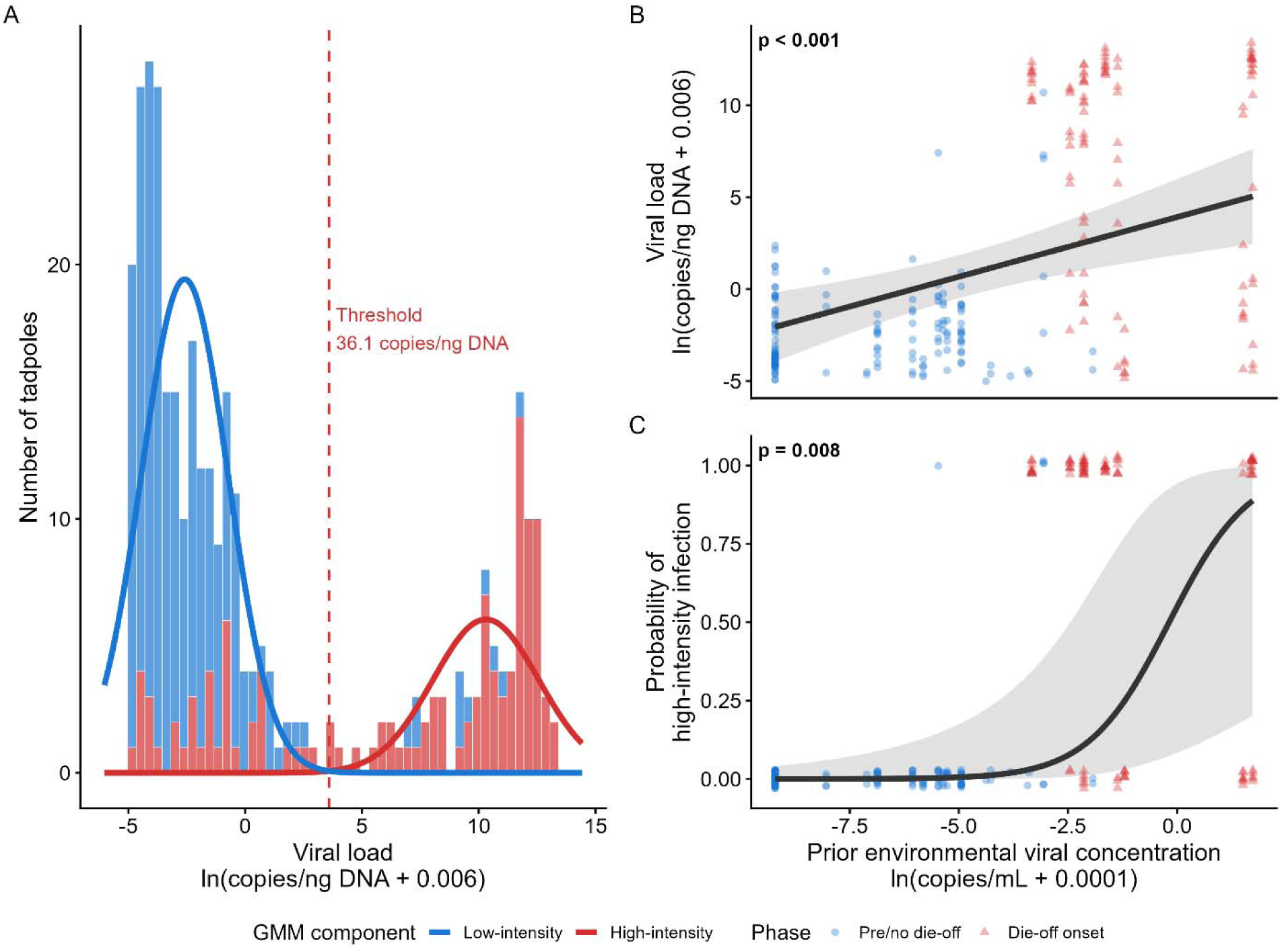
Viral load and the bimodal viral load distribution. (A) Distribution of viral loads among 337 infected tadpoles, colored by sampling context (red: die-off-onset visits; blue: pre-die-off or non-die-off visits). Gaussian mixture-model curves show the low-intensity (blue) and high-intensity (red) components, and the dashed line marks the classification threshold of 36.1 copies/ng DNA. (B) Relationship between environmental viral concentration at visit *t*−1 and viral load at visit *t*. (C) Probability of high-intensity infection as a function of environmental viral concentration at visit *t*−1, with binary observations vertically jittered. Panels B and C use the 245 infected tadpoles in the modeled sample, with lagged prevalence held at its mean among those individuals. Lines show fixed-effect predictions from the two-predictor LMM (B) and binomial GLMM (C), with random effects set to zero. The shaded bands show 95% confidence intervals. Red triangles indicate die-off-onset visits; blue circles indicate pre-die-off or non-die-off visits.

Among infected individuals sampled at visits for which prior-visit viral measurements were available (N=245 from 16 pond-years), lagged eDNA concentration and lagged prevalence jointly predicted viral load (eDNA*_t-1_* β=2.51 [95% CI: 1.17-3.84], p<0.001; prevalence*_t-1_*β=1.91 [95% CI: 0.78-3.05], p=0.002; R^2^=0.39; Fig. 4B). Higher environmental viral concentration and infection prevalence at the previous visit predicted higher viral loads among infected tadpoles at the next visit. No environmental predictor improved this model (Table S2.5), and leave-one-pond-year-out cross-validation supported out-of-sample predictive accuracy (LOOCV R^2^=0.41). Modeling viral load as a binary response (low-vs. high-intensity) showed qualitatively identical results (Fig. 4C; Table S2.3).

Unlike the effect of lagged prevalence on subsequent infection prevalence, both viral load effects were strongly attenuated after excluding onset visits (N=147 individuals, 15 pond-years). Neither lagged eDNA (β=−0.03, p=0.955) nor lagged prevalence (β=0.22, p=0.338) was significant (Table S2.6). Thus, the relationships between prior viral conditions and subsequent viral load weakened substantially when die-off-onset visits were excluded.

### Environmental viral concentration

Environmental viral concentration among visits with detectable virus (N=139 visits, 62 pond-years; includes pond-years with eDNA detection but no tissue-confirmed infection) was best predicted by its own prior concentration and the prior proportion of infected hosts (eDNA*_t-1_* β=2.70 [95% CI: 2.21-3.20], p<0.001; prevalence*_t-1_*β=1.36 [95% CI: 0.82-1.90], p<0.001; R^2^=0.63; Fig. 5A,B; Table S2.4). Leave-one-pond-year-out cross-validation here yielded the highest out-of-sample predictive accuracy across all analyses (LOOCV R^2^=0.61).

**Figure 5.**
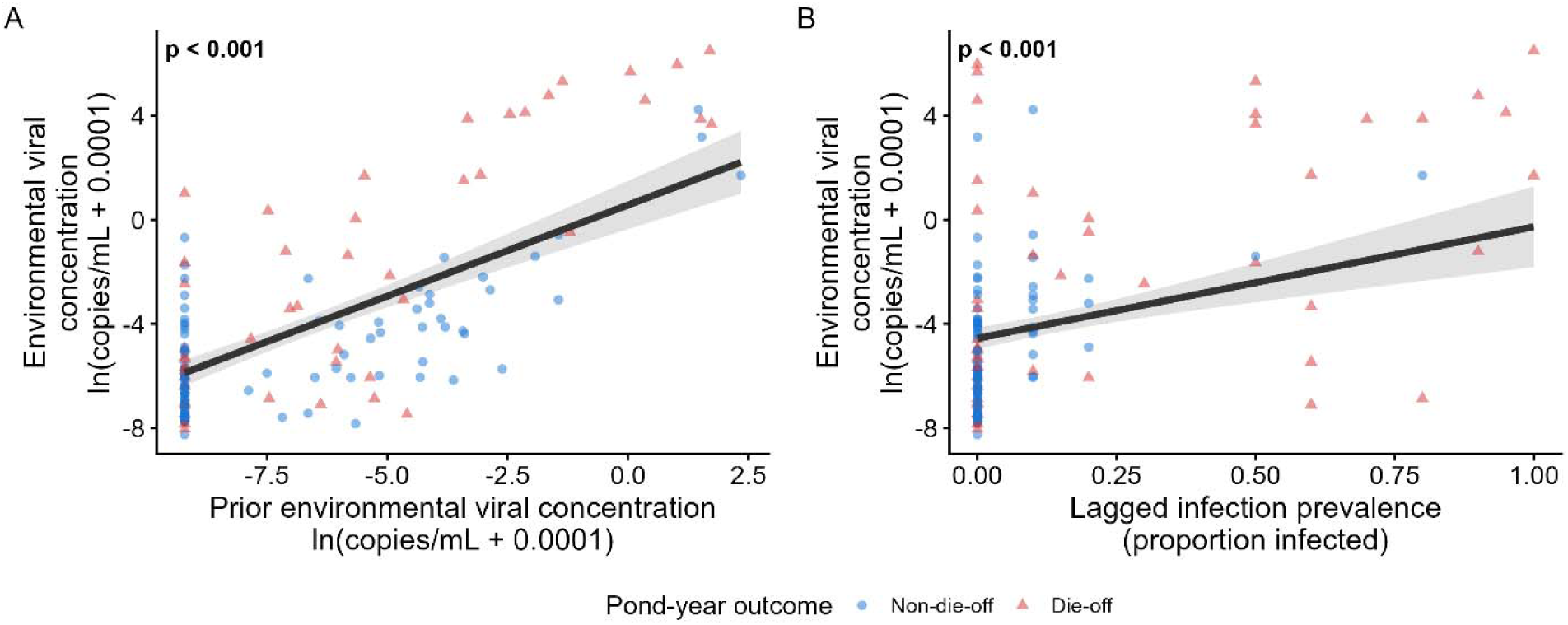
Temporal drivers of environmental viral concentration. (A) Relationship between environmental viral concentration at visit *t*−1 and environmental viral concentration at visit *t*, holding lagged infection prevalence at its mean among the modeled visits. (B) Relationship between infection prevalence at visit *t*−1 and environmental viral concentration at visit *t*, holding log-transformed lagged environmental viral concentration at its mean among the modeled visits. Both panels use the same two-predictor LMM with a pond-year random intercept, fitted to 139 eDNA-positive visits. Red triangles indicate die-off pond-years; blue circles indicate non-die-off pond-years. Lines show fixed-effect predictions with random effects set to zero; shaded bands show 95% confidence intervals. Annotated p-values correspond to the focal lagged predictor in each panel.

Unlike the relationships of environmental viral concentration with subsequent infection prevalence and viral load, the environmental viral concentration model was robust to exclusion of die-off-onset visits, and both lagged eDNA concentration and lagged prevalence remained supported (Table S2.6). Lagged eDNA concentration also remained supported when non-die-off pond-years and pre-onset visits from die-off pond-years were analyzed separately. Temperature’s association with environmental viral concentration weakened substantially after lagged viral predictors were included (β=1.36 to 0.10; p=0.609). Viral predictors also remained supported when environmental predictors were lagged. Lagged pH improved the infection prevalence model and lagged depth improved the environmental viral concentration model, but neither displaced the viral predictors (Table S2.5).

### eDNA-infection feedback

We evaluated six directional time-lagged relationships among environmental viral concentration, infection prevalence, and viral load (N=44-97 visits per test, depending on available response and lagged predictors; Table S2.7). Three of the four focal pathways linking environmental virus with infection were supported after Holm correction. Environmental viral concentration predicted subsequent infection prevalence (eDNA*_t-1_*→prevalence*_t_*: β=1.57 [95% CI: 0.76-2.38], p<0.001) and viral load (eDNA*_t-1_*→viral load*_t_*: β=2.80 [95% CI: 1.09-4.51], p=0.010), and infection prevalence predicted subsequent environmental viral concentration (prevalence*_t-1_*→eDNA*_t_*: β=1.21 [95% CI: 0.60-1.83], p=0.001). The fourth focal pathway, viral load*_t-1_*→eDNA*_t_*, was positive but not significant (β=0.98, p=0.109). Of the two additional pathways, prevalence predicted subsequent viral load, whereas viral load did not predict subsequent prevalence.

The prevalence-to-eDNA pathway, consistent with shedding from infected hosts into the water column, remained supported after excluding die-off-onset visits (β=0.95 [95% CI: 0.27-1.63], p=0.048). In contrast, the eDNA-to-prevalence and eDNA-to-viral load pathways were not supported after onset exclusion (both p=0.928; Fig. 6). Thus, during pre-die-off buildup, infection prevalence predicted subsequent environmental viral concentration. Near the transition to mass mortality, environmental viral concentration also predicted subsequent infection prevalence and viral load, completing the feedback structure shown in Fig. 6.

**Figure 6.**
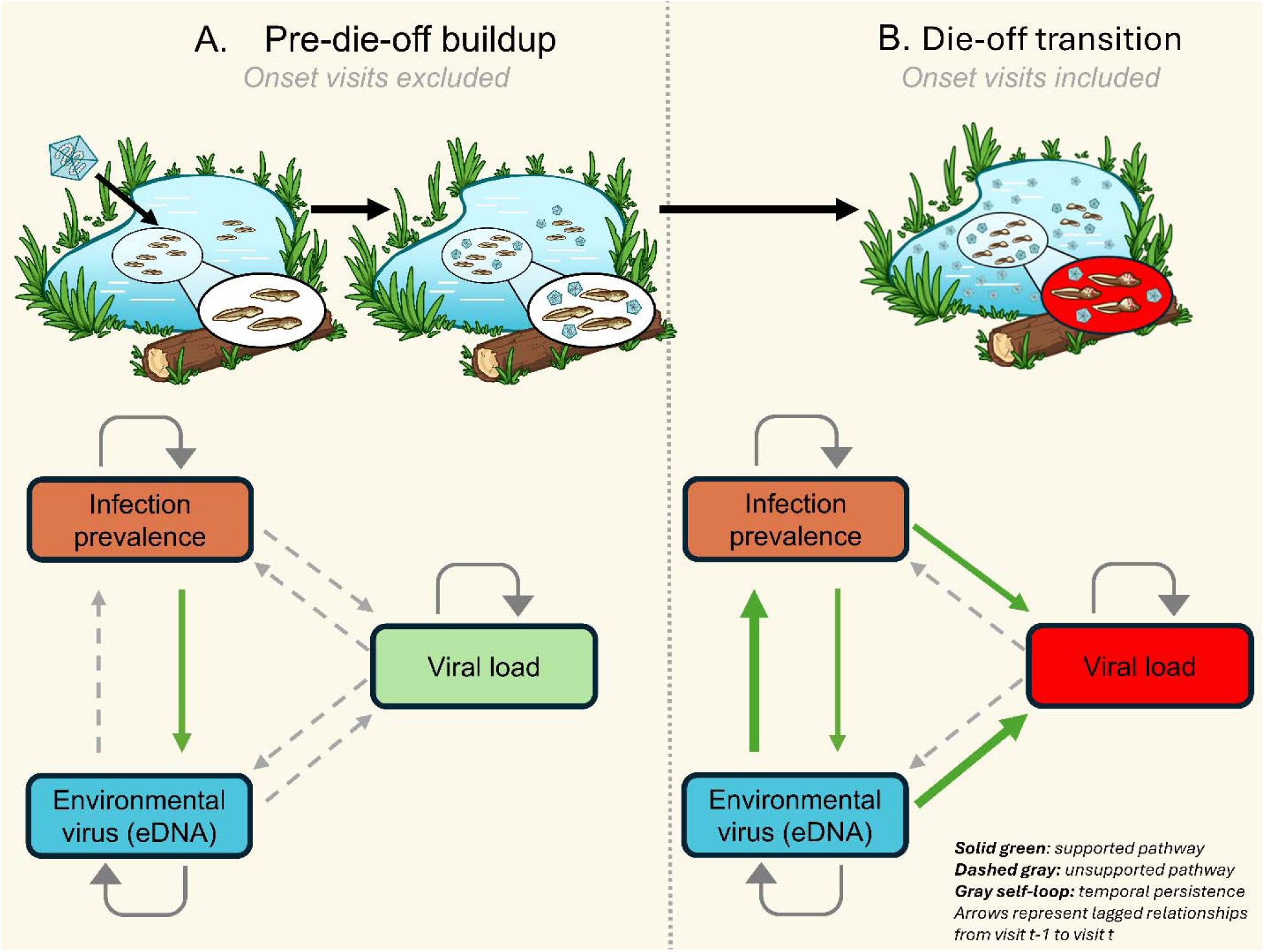
Empirical feedback structure of ranavirus epizootics. Six cross-variable arrows represent the directional time-lagged relationships evaluated among environmental viral concentration, infection prevalence, and viral load (visit t−1→visit t). Self-loops represent each variable’s temporal persistence. Solid green arrows indicate pathways supported after Holm correction across the six directional tests; dashed gray arrows indicate unsupported pathways. (A) During pre-die-off buildup, with onset visits excluded, infection prevalence predicted subsequent environmental viral concentration, whereas the other cross-variable pathways were unsupported. (B) When die-off-onset visits were included, environmental viral concentration predicted subsequent infection prevalence and viral load, infection prevalence continued to predict environmental viral concentration, and prevalence additionally predicted subsequent viral load. The two pathways originating from viral load were unsupported. Piecewise structural equation modeling provided an adequate fit when the supported pathways were estimated simultaneously (Fisher’s C=3.33, df=4, p=0.504; Table S2.8). Red shading of the viral-load state represents the shift toward high-intensity infections associated with the transition to mass mortality.

Piecewise structural equation models were consistent with the same pathway structure when prevalence, viral load, and environmental viral concentration were estimated simultaneously. The selected SEM included bidirectional eDNA-prevalence paths, eDNA*_t-1_*→viral load*_t_*, prevalence*_t-1_*→viral load*_t_*, and temporal-persistence terms for all three responses, with adequate global fit (Fisher’s C=3.33, df=4, p=0.504; N=36 visits, 15 pond-years; Table S2.8). Within this simultaneous model, the eDNA-to-prevalence path was weaker than in the pairwise model, but all estimated cross-lagged paths retained the same sign.

### Die-off discrimination and viral trajectories

Among the 32 ever-infected pond-years, no static pond feature distinguished die-off from non-die-off pond-years (all p>0.28; Appendix S2), and ponds showed little consistency in die-off occurrence across years (intraclass correlation coefficient=0.063). We also found no evidence that conditions at first detection differed between eventual die-off and non-die-off pond-years.

Trajectory slopes represented within-pond-year changes in prevalence or ln(eDNA+0.0001) over day of year. Among 14 candidate predictors, an early-season tadpole density estimate was associated with faster eDNA accumulation (β=0.086, p=0.029; N =17), while the association between earlier detection and faster prevalence increases was limited to die-off pond-years (Table S2.10).

Dynamic viral trajectories distinguished die-off from non-die-off pond-years more clearly than static features. Lagged eDNA discriminated onset visits among ever-infected pond-years with high accuracy (AUC=0.949, 95% CI [0.892–0.990]; Fig. S2.6). At the pond-year level, prevalence slope was the strongest discriminator of die-off (β=2.08 [95% CI: 0.80–4.50], p<0.001, N=28), followed by eDNA accumulation rate (β=1.51 [95% CI: 0.33–3.65], p=0.007, N=17; Table S2.9). Because die-off trajectories used only pre-onset visits, whereas non-die-off trajectories covered the full season, these comparisons describe retrospective discrimination rather than prospective prediction. The estimated lagged-eDNA cutpoint was sensitive to which pond-years were included (Fig. S2.7).

### Cross-dataset comparison

We applied parallel models to an independently collected dataset from the same forest (Hall *et al*. 2018; 8 ponds, 1 year). Viral load was available as a mean for each visit, so the viral load analysis used visits rather than individuals. Lagged eDNA and prevalence received substantially more support than seasonal and temperature predictors of mean viral load (N=25 visits; ΔAICc=13.9; viral R^2=^0.73 vs. environmental R^2^=0.53). Evidence for infection prevalence was less consistent: environmental predictors received greater support, although adding lagged eDNA produced a competitive model (ΔAICc=0.5; R^2^=0.73; Fig. S2.8). Overall, the independent analysis supported the viral load result more strongly than the infection prevalence result.

## Discussion

Ranavirus infections were widespread across our study landscape, yet die-offs remained relatively uncommon (13 of 102 pond-years; 12.7%) and were not distinguished by static pond features or conditions measured at first detection. Much of this apparent unpredictability became interpretable when we included lagged measures of the viral environment itself. Infection prevalence predicted subsequent environmental viral concentration throughout epizootics, whereas environmental viral concentration predicted subsequent infection prevalence and viral load only when die-off onset visits were included. This temporal structure is consistent with a feedback in which virus accumulates in the shared water column during a prolonged buildup and, near a die-off onset, contributes to further infection and disease escalation. A mechanistic model based on the observations in this system shows that dose-dependent progression through a shared viral reservoir can generate comparable two-phase dynamics and a sharp transition between low and high mortality without requiring among-host heterogeneity in susceptibility (Billet *et al*. 2026). Together, the field patterns and model suggest environmental viral accumulation as a plausible mechanism contributing to the occurrence and timing of die-off events.

The temporal pattern among viral states provides the clearest evidence for this feedback. Prior prevalence remained associated with subsequent prevalence during the pre-die-off buildup, as expected under temporal persistence in an ongoing epizootic. In contrast, we found that infection prevalence predicted subsequent environmental viral concentration throughout epizootics, whereas environmental viral concentration predicted subsequent host infection most strongly near die-off onset. The dose-dependent nature of ranavirus pathogenesis and the limited acquired immunity of larval amphibians provide a potential biological basis for this pattern (Andino *et al*. 2012; Brunner *et al*. 2005; Rollins-Smith 1998; Warne *et al*. 2011): rising environmental viral concentrations in a bounded water body can increase the dose experienced by many hosts simultaneously, linking new infections and within-host escalation through the same environmental reservoir. New infections and within-host disease progression therefore remain biologically distinct processes, but may be coupled through the same changing exposure environment. Whether accumulation proceeds far enough to produce this escalation should depend on the balance between viral inputs and losses (host density and shedding, water volume and renewal, viral decay, microbial degradation) as well as host susceptibility. In managed systems, interventions that increase water exchange, reduce host crowding, or accelerate pathogen inactivation could slow environmental accumulation even after infection has established. The shared environment may therefore function not only as a source of infection but as a reservoir whose changing viral concentration influences population-level disease escalation.

A potential alternative interpretation of the relationship between environmental viral concentration and viral load is that observed viral loads rose near die-off because highly infected individuals died before they could be sampled at earlier visits, or because older and more developed larvae tolerated higher loads, rather than because increasing environmental viral concentration contributed to higher viral loads. Our cross-sectional sampling cannot fully distinguish these mechanisms. However, a concentration-dependent escalation interpretation is supported by the temporal ordering of the relationship, with environmental viral concentration at the preceding visit predicting viral load among infected hosts at the subsequent visit. This relationship remained after accounting for prior mean viral load and seasonal timing and was little changed by adding developmental stage. The same eDNA-viral load relationship held when we applied our analytical framework to the Hall *et al*. (2018) dataset, in which viral-state predictors were more strongly supported than seasonal and temperature predictors of mean viral load. These results support environmental exposure as a contributor to higher viral loads, while selective mortality and developmental tolerance remain plausible contributors to the observed rise in viral load.

The features associated with ranavirus occurrence did not distinguish infected pond-years that subsequently experienced die-offs. Among 14 candidate predictors of viral trajectory, tadpole density at first detection was associated with faster eDNA accumulation. Although based on a limited number of infected pond-years, this result suggests that host abundance may influence how rapidly virus accumulates after establishment. Environmental processes could affect the same trajectory. For example, seasonal water loss may increase viral concentration without changing host susceptibility. Rates of change in environmental viral concentration and prevalence distinguished die-off from non-die-off pond-years more clearly than conditions at first detection or viral state at a single time point. This pattern is consistent with a tipping-point process in which gradual viral accumulation leads to an abrupt transition (Scheffer *et al*. 2009). Our field data did not, however, identify a clear transition threshold. Transition points varied among pond-years, likely because local conditions altered the balance between viral inputs and losses and because biweekly sampling cannot precisely resolve the rapid escalation often observed. Thus, the distinguishing feature of die-off pond-years appears not to be a consistent starting condition but a faster viral trajectory that culminates in phase-dependent feedback near mortality onset.

These results do not imply that host susceptibility or abiotic conditions are unimportant factors in mass mortality. Indeed, wood frogs are among the North American amphibian species most susceptible to ranavirus, which undoubtedly contributes to their frequent involvement in reported die-off events (Hoverman *et al*. 2011). Developmental stage contributed beyond the viral predictors for infection prevalence, indicating that host ontogeny remains an important component of infection dynamics not fully captured by our framework (Billet & Skelly 2025). Temperature can also influence host defenses, viral replication, and environmental persistence (Brand *et al*. 2016; Nazir *et al*. 2012; Rollins-Smith 2017). However, temperature provided little independent predictive information once recent viral conditions were included. Whether this attenuation reflects effects mediated through viral dynamics or confounding by shared seasonality cannot be cleanly resolved here. More broadly, factors that alter individual susceptibility may shape host-level outcomes without necessarily determining whether pathogen accumulation develops into a population-level process. This distinction may help explain why experimental manipulations of susceptibility have often produced weak or inconsistent changes in ranavirus epizootic outcomes (Brunner *et al*. 2025a; Haislip *et al*. 2012; Reeve *et al*. 2013).

Our mechanistic model predicts that environmental viral accumulation depends on the host-to-water-volume ratio and can cross the progression threshold rapidly when that ratio is high (Billet *et al*. 2026). In tanks and mesocosms, limited dilution may therefore compress differences among susceptibility treatments, making experimental venue itself part of the scaling from individual responses to population outcomes (Skelly 2002).

The mechanism suggested here may extend beyond larval amphibians wherever infected hosts shed pathogen into a bounded shared environment and disease severity increases with exposure. Ranavirus die-offs in the United Kingdom affect adult common frogs in small garden ponds rather than larvae in vernal pools (Campbell *et al*. 2018; Teacher *et al*. 2010). Despite substantial differences in host life stage, body size, and immune capacity, limited viral dilution in small, enclosed water bodies is a common feature. Similar reservoir-mediated dynamics have been proposed in Bd-amphibian systems, where repeated exposure to a shared zoospore pool can elevate infection intensity beyond levels associated with population decline (Briggs *et al*. 2010; Vredenburg *et al*. 2010), and in OsHV-1 oyster systems, where increased water renewal reduces viral accumulation and mortality (Petton *et al*. 2015). Theoretical models likewise show that shared environmental reservoirs can generate threshold-dependent dynamics distinct from those represented by simple direct-transmission models (Codeço 2001; Tien & Earn 2010). These examples do not establish that the same feedback operates across systems, but they point to a broader ecological principle: in shared environments with limited exchange, population outcomes may depend on the balance between pathogen inputs from shedding and losses through decay, dilution, and removal, rather than on individual susceptibility alone.

Our study encompasses 13 die-off events across 102 pond-years at a single site involving a single host species. Whether the hypothesized feedback structure generalizes across sites, host taxa, and viral strains remains to be tested. Despite weekly mortality surveys, brief or low-magnitude events may have gone undetected; the 13 observed die-offs should therefore be considered a minimum (Brunner & Calabrese 2025). Further, the single-forest design constrains abiotic variation to some degree (Cohen *et al*. 2016), though ponds do vary substantially in canopy cover, morphology, temperature, and proximity to road runoff (Fig. S2.2). Additionally, directional time-lagged analyses establish temporal ordering but not definitive causation, and environmental viral concentration could covary with unmeasured host, pathogen, or environmental factors that independently influence infection outcomes (Billet *et al*. 2025; Brenes *et al*. 2014; Brunner *et al*. 2017; Savage *et al*. 2019; Wuerthner *et al*. 2017). Moreover, eDNA indexes environmental viral genetic material, including degraded or noninfectious DNA, rather than infectious virions or exposure dose. Experimental manipulation of infectious viral concentration, water renewal, or host-to-water-volume ratio, coupled with direct measures of viral infectivity, would help determine whether environmental viral accumulation contributes causally to progression to high-intensity infection or primarily tracks it. Finally, the association between density at first detection and eDNA accumulation rate rests on a small sample of infected pond-years (N=17) and may interact with unmeasured factors such as initial inoculum size or stochastic early infection dynamics.

Across systems, severe outcomes may depend not simply on pathogen presence but on how pathogen burden and exposure change through time (Briggs *et al*. 2010; Bruno *et al*. 2007; Kock *et al*. 2018; Petton *et al*. 2015). Our results suggest a plausible mechanism by which such dynamics can arise: in physically bounded environments, a shared pathogen reservoir can integrate shedding across hosts and alter the exposure experienced by the population. In our system, trajectories of environmental viral concentration and infection prevalence were more informative than static pond conditions or infection status at a single point in time, although the field data do not establish that reservoir dynamics alone determine whether or when die-offs occur. More broadly, host susceptibility may shape individual infection outcomes, but whether infection escalates to population-level mortality may depend on collective pathogen accumulation in the shared environment. Wastewater-based epidemiology similarly illustrates how environmental sampling can integrate shedding across hosts and reveal population-level pathogen dynamics not readily apparent from individual observations (Peccia *et al*. 2020).

Surveillance that tracks environmental pathogen trajectories alongside host infection and habitat conditions may therefore improve detection of escalating outbreaks and make outcomes that have long appeared stochastic more predictable.

## Acknowledgments

We thank Adriana Rubinstein, L. Kealoha Freidenburg, Carolyn Skotz, Yara Alshwairikh, Sydney Nelson, and Karinne Tennenbaum for their assistance with fieldwork and/or labwork. We thank Trenton Garner, David Klinges, and Andis Arietta for useful suggestions and conversations around this work. Animal handling was approved by Yale University IACUC protocol 2019-10361 and 2022-10361. Specimens were collected under CT DEEP Permit 0124002. Field surveys were conducted with permission from Yale Myers Research Committee (BILL21).

