## Appendix 1 for "Epizootic tipping points: Environmental viral feedbacks predict amphibian die-offs"

Supplementary materials for: Epizootic tipping points: Environmental viral feedbacks predict amphibian die-offs

### Appendix S1: Supplementary methods

#### S1.1 Field sampling protocols

Wood frogs typically bred in late March to early April at YMF. We censused each pond for egg masses within approximately one week of the first observed breeding activity each spring, estimating total counts as the average of two independent counts. Egg mass counts serve as an reasonable proxy for breeding female abundance (Berven & Grudzien 1990; Crouch & Paton 2000).

Ponds were visited at approximately biweekly intervals from April through July (~630 total visits across the study). At each visit, we conducted timed dip-net surveys scaled to pond surface area to estimate tadpole density, following Werner *et al.* (2007). We haphazardly collected tadpoles with a target of 20 individuals per visit, collecting fewer when breeding effort was low to prevent overcollection or when 20 tadpoles were not captured during the survey. We recorded conductivity, pH, dissolved oxygen, and maximum pond depth at each visit.

At each visit, we collected environmental DNA samples by pooling ten 100-mL surface water samples from locations distributed across the pond perimeter into sterile Whirl-Pak bags. Samples were stored on ice and transported to the laboratory for vacuum filtration within 24 hours of collection.

Pond water temperatures were recorded continuously at 30-minute intervals using HOBO pendant loggers (Onset Computer Corporation, Bourne, MA, USA). Approximately 30% of records were missing due to logger loss; missing daily temperatures were imputed using Random Forest models (R^2^ = 0.95, RMSE = 1.0°C; see Billet & Skelly 2025, Appendix S1 for full methodological details). From the complete daily series, we calculated visit-level 7-day rolling means and standard deviations, season-level metrics (May 1-June 30), and temperature anomalies (residuals of 7-day mean temperature regressed on day of year).

Ponds were visited weekly (more frequently than biweekly sampling visits) for visual encounter surveys of carcasses. Die-off onset was defined as the first visit at which five or more carcasses were detected, following Hall *et al.* (2018). Surveys typically covered the entire pond perimeter and accessible shoreline, with consistent effort across visits. All equipment (dip nets, waders, sampling tools) was sanitized between sites using 2% chlorhexidine diacetate solution (Bryan *et al.* 2009).

Static pond attributes included canopy openness (weighted global site factor from hemispherical photography; Arietta *et al.* 2020), surface area (ellipse approximation), elevation (elevatr package; Hollister *et al.* 2023), and surrounding land cover (NLCD within 200-m buffer; Homer *et al.* (2012); see Billet *et al.* (2024) for details). Missing canopy values (2 ponds) were imputed with the median (GSF = 20.1).

#### S1.2 Laboratory methods

##### S1.2.1 Tissue processing, DNA extraction, and quantitative PCR

Collected tadpoles were euthanized in 10% ethanol in the field (Beaupre *et al.* 2004) and preserved in 95% ethanol. Tadpoles were never cohoused while alive. Tadpoles were staged following Gosner (1960), measured (snout-vent length), and dissected to obtain liver tissue. DNA was extracted using Qiagen DNeasy Blood & Tissue kits (Qiagen, Hilden, Germany) following the manufacturer's protocol. Extraction negative controls were included to monitor for contamination. DNA concentration in extractions was measured using a NanoDrop 2000c (ThermoFisher Scientific, Waltham, MA, USA).

We performed qPCR using primers and a FAM-labeled probe targeting a 97-bp region of the ranavirus major capsid protein (MCP) gene (Leung *et al.* 2017). Each 20-µL reaction contained 2 µL template DNA, 1 µL each of forward and reverse primers (10 µM), 0.05 µL probe (100 µM), 10.0 µL SsoAdvanced Universal Probes Supermix (Bio-Rad Laboratories, Hercules, CA, USA), and 5.95 µL Nanopure water. Each qPCR plate included a standard dilution series (2.79 × 10^6^ - 2.79 × 10^1^ copies) prepared from a 350-bp synthetic gBlock (IDT, Coralville, IA, USA) containing the MCP target region, plus a negative control of Nanopure water. Each standard, control, and unknown sample was run in duplicate on 96-well plates using a CFX Connect thermocycler (Bio-Rad Laboratories). Cycling conditions were 98°C for 3 min followed by 40 cycles of 98°C for 15 s and 60°C for 45 s, with a plate read at the end of each cycle. Samples for which both duplicate reactions amplified before cycle 40 were considered positive. When duplicates were discrepant, a third reaction was run. If the third reaction amplified, the sample was considered positive. Viral quantities were averaged across positive duplicate wells and expressed as viral copies per ng DNA using NanoDrop-derived DNA concentrations. No plate negative or extraction negative controls amplified.

Among 220 positive tissue samples with viral loads below 1 copy/ng DNA, 18 (8.2%) had estimated quantities below one copy per reaction (range: 0.25-0.99 copies). Each positive classification was supported by amplification in at least two reactions. Excluding these detections did not change the best-supported continuous viral-load model or its conclusions.

##### S1.2.2 eDNA filtration, DNA extraction, and quantitative PCR

Water samples were vacuum-filtered within 24 hours of collection through 0.45-µm nitrocellulose filters (Nalgene Analytical Filter Funnels, ThermoFisher, Waltham, MA, USA) using a powered vacuum pump until all water was filtered or the filter clogged with debris (mean volume filtered: 666 mL; range: 100-1,100 mL). Filters were folded using forceps cleaned in 50% bleach and placed in 2-mL tubes filled with 95% ethanol. Laboratory process blanks were prepared by placing 1,000 mL of deionized water in the same sterile Whirl-Pak bags used for field samples and carrying the water through the same filtration, extraction, and qPCR process. Preserved filters were stored at -20°C until processing. DNA was extracted from filters using the QiaShredder and DNeasy Blood & Tissue kit method (Goldberg *et al.* 2011), with extraction negative controls included.

For eDNA samples, we used the same primer-probe set and general plate setup as for tissue samples. Each 20-µL reaction contained 2 µL template DNA, 1 µL each of forward and reverse primers (10 µM), 0.05 µL probe (100 µM), 10.0 µL TaqMan Environmental Master Mix 2.0 (Applied Biosystems), and 5.95 µL Nanopure water. One duplicate well per eDNA sample was spiked with TaqMan Exogenous Internal Positive Control (ThermoFisher Scientific, Waltham, MA, USA) to test for PCR inhibition. Cycling conditions were 50°C for 5 min, 95°C for 10 min, then 40 cycles of 95°C for 15 s and 60°C for 60 s, with a plate read at the end of each cycle. Environmental viral concentration (eDNA) was calculated as viral copies per mL of water filtered. No samples showed evidence of inhibition, and no laboratory process blanks, extraction negatives, or plate negatives amplified.

#### S1.3 Data processing and analysis

Tadpole density estimates were missing for approximately 32% of visits, predominantly early-season visits when tadpoles were too small or not yet dispersed from the oviposition site. For wet visits before day of year 182, missing density was assigned the earliest available measurement from the same pond-year. Missing density values at later and dry-pond visits remained missing. For tissue-confirmed occurrence analyses, pond-year median density was calculated from non-die-off or pre-die-off visits occurring before day of year 182 to reduce bias from mortality and metamorphosis.

Remaining occasional missing values in visit-level water chemistry (pH, conductivity, dissolved oxygen, and depth) and temperature summaries were imputed using the median for that pond-year for wet visits; values from dry visits remained missing. Missing pond-year canopy openness and seasonal temperature summaries were imputed using study-wide medians.

Of 7,955 collected tadpoles, 2,685 (33.8%) were screened for ranavirus infection via qPCR. Screening effort was higher in die-off pond-years (~48% of collected individuals screened) than in non-die-off pond-years (~26%). Screening was allocated to focal visits within each pond-year, with additional effort during die-offs to characterize rapid changes in infection prevalence and viral load. Within each focal visit, individuals selected for qPCR were chosen at random from the collected pool. Because not every visit included tissue screening, lagged prevalence or viral load was sometimes missing when the preceding visit had no screened individuals. These observations were excluded by complete-case filtering and screening-gap lags were not imputed. To assess whether unequal screening effort affected the infection prevalence results, we conducted 500 random-subsampling iterations in which tissue screening was capped at 10 randomly selected individuals per visit. In each iteration, we recomputed visit-level prevalence and its one-visit lag, retained visits with tissue screening at both the current and immediately preceding visits, and refit the primary infection prevalence GLMM using the same predictor scaling and random-effects structure. For the first visit in each pond-year, lagged prevalence was set to zero. Log-transformed lagged eDNA and mean viral load were set to ln(0 + 0.0001) and ln(0 + 0.006), respectively.

Leave-one-pond-year-out cross-validation (LOOCV) iteratively withheld all observations from a single pond-year, fit the model to the remaining data using fixed-effects predictions only, and evaluated predictive accuracy on the held-out observations. Predictive R^2^ was computed as 1 minus the ratio of residual to total sum of squares. Pond-year was the appropriate cross-validation unit because individuals and visits within a pond-year share unmeasured environmental conditions and disease trajectories.

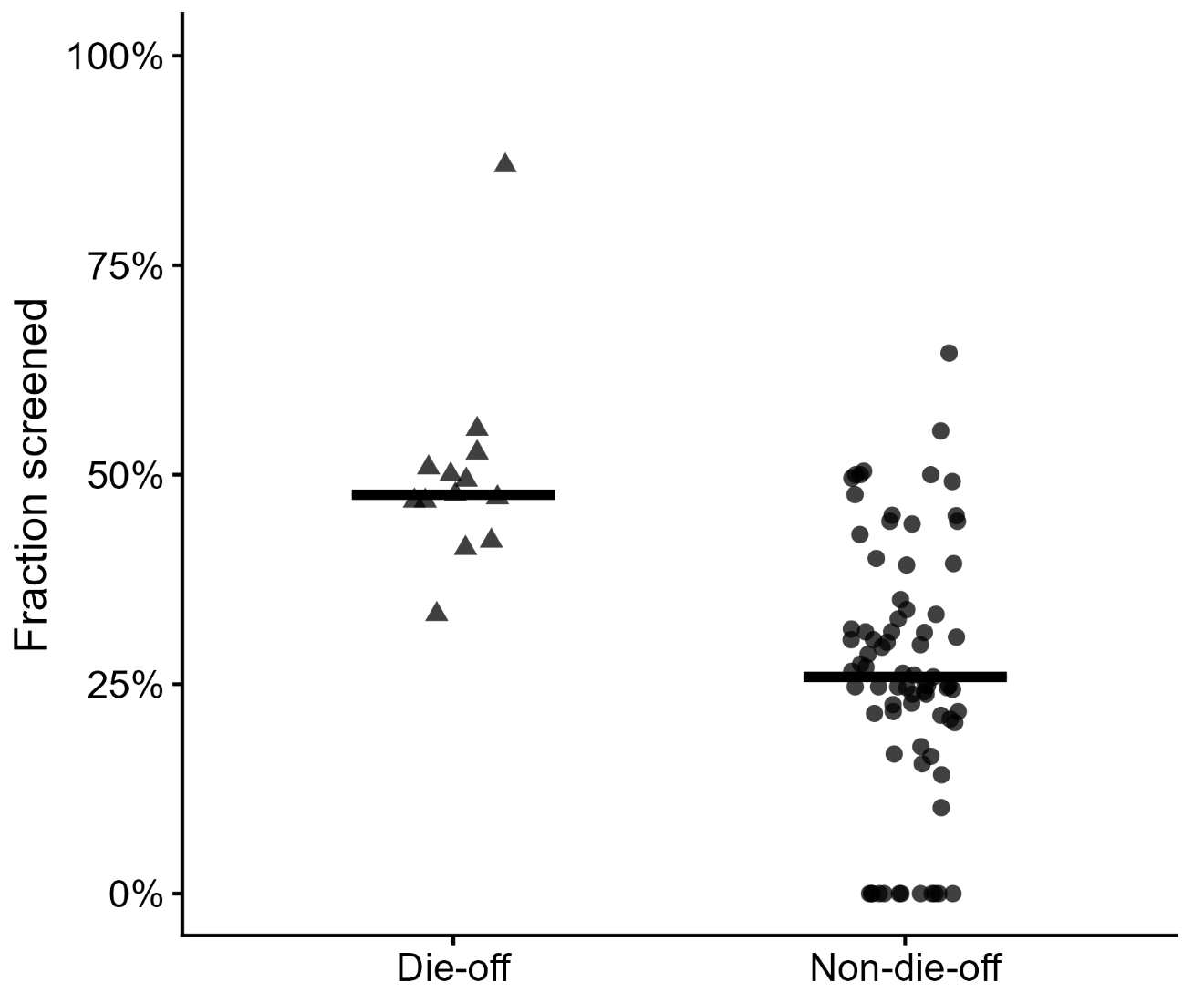

**Figure S1.1.** Screening effort by pond-year and fate. Points represent 90 pond-years with collected tadpoles (13 die-off and 77 non-die-off), using visits through die-off onset and excluding post-onset visits. The y-axis shows the fraction of collected individuals screened by qPCR. Triangles indicate die-off pond-years and circles indicate non-die-off pond-years; horizontal lines show group medians (approximately 48% and 26%, respectively).

#### S1.4 Extended statistical analysis

##### Software and computation

Mixed-effects models were fit using lme4 (Bates *et al.* 2015). Model diagnostics used DHARMa scaled residuals (Hartig *et al.* 2024) with 1,000 simulations per model. Structural equation models were fit using piecewiseSEM (Lefcheck 2016). Firth penalized logistic regression used the logistf package (Heinze & Schemper 2002), and ROC analysis used pROC (Robin *et al.* 2011). AICc was computed via MuMIn (Bartoń 2023), and spatial autocorrelation tests used spdep (Bivand *et al.* 2026). Figures were produced with ggplot2 (Wickham 2016). Marginal R^2^ (variance explained by fixed effects only) and conditional R^2^ (fixed plus random effects) were computed via MuMIn::r.squaredGLMM() using the default theoretical/delta method.

##### Variable transformations

All log transformations in our study used the natural logarithm unless otherwise noted. Individual viral load (copies per ng DNA) was transformed as ln(x + 0.006). eDNA concentration (copies per mL) was transformed as ln(x + 0.0001). All of our continuous predictors were centered and scaled across the full dataset prior to subsetting for specific analyses, with the same scaling parameters retained across sensitivity analyses.

Two predictors were residualized against day of year (DOY) to remove seasonal covariation. Temperature residuals were obtained from a linear regression of 7-day mean temperature on DOY fit across all visit-level observations, capturing within-season temperature anomalies relative to the seasonal warming trend. Developmental stage residuals were obtained from a linear regression of Gosner stage on DOY fit across all individual-level observations, capturing developmental advancement or delay relative to the population-average trajectory. Visit-level mean stage residuals were computed as the mean of individual stage residuals per visit.

##### Bimodality of viral load

Bimodality of individual viral loads among infected tadpoles was assessed using two complementary approaches. Hartigan's dip test (diptest package; Maechler & Ringach 2025) tested the null hypothesis of unimodality against a multimodal alternative. A two-component Gaussian mixture model (GMM) was fit to ln-transformed viral loads using the expectation-maximization algorithm (normalmixEM, mixtools package; Young *et al.* 2025) with k = 2 components, convergence tolerance ε = 10^-8^, and maximum 1,000 iterations. Components were ordered by mean (lower = low-intensity, higher = high-intensity). The classification threshold was computed as the crossing point where the posterior probabilities of the two components are equal. We used the GMM-derived crossing point as the threshold for binary viral load analyses.

##### Common-sample enforcement

Within each model group, all Phase 0 and Phase 1 candidate models were fit on an identical complete-case dataset for fair AICc comparison. The common sample required non-missing values for every predictor included in the Phase 0-1 comparison set (Table S1.1). When testing individual environmental additions to the best viral model (Phase 2), each predictor’s ΔAICc was computed on that predictor’s complete-case subset with the base model refit on the same data because some Phase 2 predictors had additional missing values (e.g., pH and dissolved oxygen).

##### Model diagnostics

Residual diagnostics used DHARMa scaled residuals with 1,000 conditional simulations per model. We assessed residual uniformity and dispersion, examined QQ and residual-versus-predicted plots, and checked for temporal autocorrelation. Collinearity was assessed using variance inflation factors (VIF; Fox & Weisberg 2019).

The diagnostic tests showed departures from residual uniformity for the prevalence and viral load models and fitted-value-dependent patterns for prevalence and environmental viral concentration (Figure S2.9). Viral load quantile tests were not interpreted because the fits produced convergence warnings. No comparable departures were detected for the binary high-intensity model. The focal associations were robust to the binary low-/high-intensity formulation of viral load and to pond-year-clustered CR2 inference for environmental viral concentration.

##### Confidence interval computation

We report 95% Wald confidence intervals (β ± 1.96 × SE) for all mixed-effects model coefficients (GLMMs and LMMs fitted via lme4). For Firth penalized logistic regression models, we report profile-likelihood confidence intervals. Bootstrap confidence intervals (2,000 replicates, percentile method, resampling pond-years with replacement) were used to estimate ROC AUC.

##### Three-phase model comparison details

Phase 0 (nonviral) and Phase 1 (viral-only) candidate sets are enumerated in the AICc tables for each analysis (Tables S2.1-S2.4); all candidates within a family were compared on a common sample. Phase 2 tested individual environmental predictors as additions to the best viral model; the specific predictor sets varied by analysis and are listed in Table S2.5. For eDNA detection, the "lagged eDNA" predictor was binary (detected vs. not detected at the prior visit) rather than the continuous log-concentration used in other analyses.

To test whether using concurrent environmental conditions disadvantaged environmental predictors relative to the lagged viral predictors, we repeated the Phase 2 analyses using temperature, depth, conductivity, and pH measured at the preceding visit. Each lagged environmental predictor was added individually to the viral base model, with the base model refitted to the same complete-case subset.

To test sensitivity to temperature parameterization, we tested raw 7-day mean temperature and the day-of-year-residualized temperature anomaly separately as Phase 2 additions to each viral base model, refitting the base model on the corresponding complete-case subset.

##### Directional time-lagged tests

Six directional tests evaluated all pairwise temporal relationships among eDNA concentration, infection prevalence, and mean viral load. Each model included the response variable’s own one-visit lag and day of year to account for temporal persistence and seasonal trends. We report cross-lagged coefficients with 95% Wald confidence intervals (β ± 1.96 × SE); the six Wald p-values were adjusted together using the Holm method to control the family-wise error rate. Sample sizes differed among tests because each required complete data for its specific response, autoregressive predictor, and cross-lagged predictor (Table S2.7). For environmental viral concentration, we additionally compared the temperature coefficient from a temperature-only model with that from a model including lagged environmental viral concentration and infection prevalence.

##### Piecewise structural equation modeling

We used piecewise SEM (Lefcheck 2016) to simultaneously estimate the directional pathways among prevalence, mean viral load, and environmental viral concentration.

The SEM required complete data for all three responses and all three lagged predictors at each visit, yielding N = 36 visits from 15 pond-years. The main limiting predictor was lagged mean viral load, which requires the previous visit to have had screened, infected individuals. Because viral load entered the SEM as visit-level means rather than individual observations, the viral load equation used (1|pond_year) without the visit-level random intercept used in the main viral models at the individual level. DOY was excluded from all SEM components because cross-validation indicated poor out-of-sample predictive accuracy and, with OLRE, it was non-significant in all three equations. All SEM models included correlated error terms among contemporaneous responses (prevalence-viral load, prevalence-eDNA, viral load-eDNA) to account for unmeasured shared drivers at each visit.

We compared three constrained pathway models and selected the simplest model with adequate overall fit (Fisher’s C, p ≥ 0.05). Model A included three temporal-persistence paths, bidirectional eDNA–prevalence paths, and a path from lagged eDNA to viral load. Model B added a path from lagged viral load to eDNA, whereas Model C added a path from lagged prevalence to viral load. A saturated model including all cross-lagged paths was retained as a fallback but was not required. Onset sensitivity was assessed by refitting the selected model after excluding die-off-onset visits (N = 36 to N = 27).

##### Ranavirus occurrence models

Ranavirus occurrence was modeled at the pond-year level using two responses: tissue-confirmed occurrence (at least one tissue-positive individual) and eDNA occurrence (two or more positive eDNA samples). For each response, we compared univariate binomial GLMMs for conductivity, egg mass count, tadpole density, area, depth, elevation, canopy openness, season mean temperature, season temperature SD, pH, wetland cover, and a null model, using a common complete-case sample of 97 pond-years from 38 ponds. The final tissue-occurrence model included conductivity and egg mass count, whereas the eDNA-occurrence model retained conductivity alone; both were refit to all 102 pond-years.

Spatial autocorrelation was tested using Moran’s I with inverse-distance weighting. A fully connected weight matrix was constructed from pond coordinates, with weights inversely proportional to Euclidean distance and row-standardized. Three tests were conducted: tissue-positive status across all ponds, die-off occurrence among ever-infected ponds, and Pearson residuals from the best tissue occurrence model. Distance to the nearest die-off pond was also tested as a fixed effect.

##### Die-off discrimination analyses

*Intraclass correlation.* Before fitting die-off discrimination models, we assessed the repeatability of die-off occurrence across years across all pond-years using intraclass correlation coefficients (ICC) from null binomial GLMMs with (1|pond) random intercepts, computed on the logistic scale as σ^2^_pond_ / (σ^2^_pond_ + π^2^/3).

*Static predictors.* Five static pond-year features (area, median tadpole density, egg count, elevation, conductivity) were tested as univariate Firth predictors of die-off among ever-infected pond-years.

*Trajectory predictors.* Five trajectory metrics were computed from pre-onset visits (for die-off pond-years) or the full season (for non-die-off pond-years): eDNA slope (OLS slope of ln(eDNA + 0.0001) versus DOY on eDNA-positive visits, minimum 3 visits), prevalence slope (OLS slope of prevalence versus DOY on screened visits, minimum 3 visits), peak prevalence (maximum visit-level prevalence), cumulative eDNA (trapezoidal AUC of ln_edna over DOY, minimum 2 visits), and eDNA peak (maximum raw eDNA concentration). For die-off pond-years, trajectory metrics were calculated using data strictly before mortality onset, whereas for non-die-off pond-years, they used the full season. This difference may make the groups appear more distinct, so the analysis shows how their trajectories differed in hindsight rather than how well die-offs could be predicted in advance.

*Trajectory determinants.* To test whether any measured variable predicted viral trajectory itself (rather than using trajectory to predict die-off, as in the Firth regression framework above), we modeled prevalence slope (N = 28 ever-infected pond-years with ≥3 tissue-screened visits) and eDNA slope (N = 17 pond-years with ≥3 eDNA-positive visits) as continuous responses in univariate OLS regressions against 14 candidate predictors (7 static pond features, 5 first-detection conditions, 2 seasonal temperature metrics; Table S2.10). For predictors reaching p < 0.05, we examined Pearson correlations separately within die-off and non-die-off pond-years as a diagnostic of whether the overall association was consistent across outcomes or concentrated within one group. Missing early-season density values were filled using the first available measurement from the same pond-year. In the eDNA-slope analysis, 14 of 17 values came from later visits, up to 29 days later, all before die-off onset.

*ROC analysis.* ROC curves assessed discrimination of die-off onset at visit *t* from environmental viral concentration measured at the preceding visit (*t*-1). Analyses were conducted for all pond-years, ever-infected pond-years (primary comparison), and die-off pond-years only (onset timing within eventual die-off pond-years). Cluster-bootstrapped 95% confidence intervals were calculated for all three analyses using 2,000 replicates with pond-years resampled with replacement and the percentile method. Threshold stability for the primary ever-infected analysis was assessed from the bootstrap distribution and 95% percentile interval of Youden’s J optimal cutpoint.

#### *Hall et al. (2018) cross-dataset comparison*

We reanalyzed an independently collected dataset from the same forest (Hall *et al.* 2018) comprising 8 ponds (4 die-off, 4 non-die-off) sampled in 2014. Hall titer values were analyzed on the scale used in the Hall dataset, without applying the natural-log transformation used for our main-study data. Lagged predictors (eDNA, prevalence, tissue titer) were constructed as one-step lags within each pond, ordered by day of year. Die-off-onset visits were retained and post-onset visits excluded, consistent with our primary analyses.

Infection prevalence was modeled with twelve candidate binomial GLMMs with pond and observation-level random intercepts. Mean viral load was modeled with thirteen candidate Gaussian LMMs with a pond random intercept, including a model with lagged tissue titer. Candidate models represented nonviral, viral-state, and combined predictors and used a common analytical sample within each response.

**Table S1.1.** Common sample sizes for Phase 0-1 model comparisons. All candidate models within each family were fit on the same complete-case dataset for AICc comparison. PY = pond-year.

| **Analysis** | **N (unit)** |
| --- | --- |
| Infection prevalence | 95 visits, 25 PYs |
| Viral load | 245 individuals, 42 visits, 16 PYs |
| eDNA detection | 130 visits, 31 PYs |
| eDNA quantity | 139 visits, 62 PYs |

**Table S1.2.** Random effects structure for each model family. OLRE = observation-level random effect for overdispersion in binomial models (Harrison 2014). Viral load models include a visit-level random intercept to account for non-independence among individuals collected at the same visit. Piecewise SEM viral load equations model visit-level means and so omit the visit-level intercept.

| **Model family** | **Random effects** | **Details** |
| --- | --- | --- |
| Pond-year tissue occurrence and eDNA occurrence | (1\|pond) | Pond-year is observation unit; ponds were sampled multiple years |
| Infection prevalence | (1\|pond_year) + (1\|obs_id) | Pond-year clusters visits; OLRE for overdispersion (Harrison 2014) |
| Viral load (continuous and binary) | (1\|pond_year) + (1\|visit_id) | Pond-year clusters visits; visit clusters individuals |
| Viral eDNA detection | (1\|pond_year) + (1\|obs_id) | Same as infection prevalence |
| Viral eDNA concentration | (1\|pond_year) | Gaussian |
| Cross-lagged (binomial) | (1\|pond_year) + (1\|obs_id) | Same as infection prevalence |
| Cross-lagged (Gaussian) | (1\|pond_year) | Same as viral eDNA concentration |
| Piecewise SEM (binomial) | (1\|pond_year) + (1\|obs_id) | OLRE for Type I error control |
| Piecewise SEM (Gaussian) | (1\|pond_year) | Visit-level means; no individual nesting |

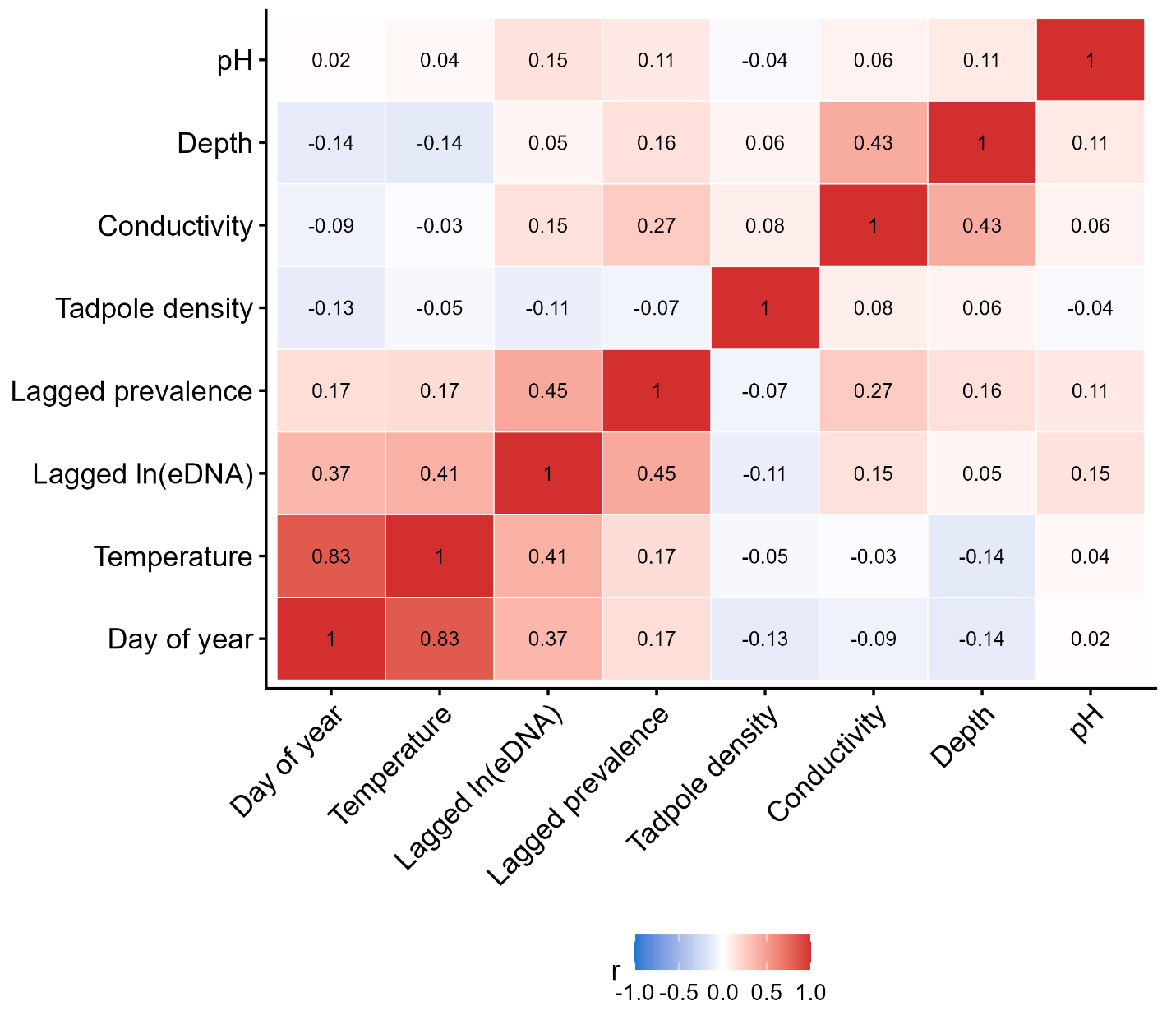

**Figure S1.2**. Predictor collinearity matrix. Pairwise Pearson correlations among eight candidate predictors, calculated from 286 visits with complete data for all eight predictors.

### Appendix S2: Supplementary results

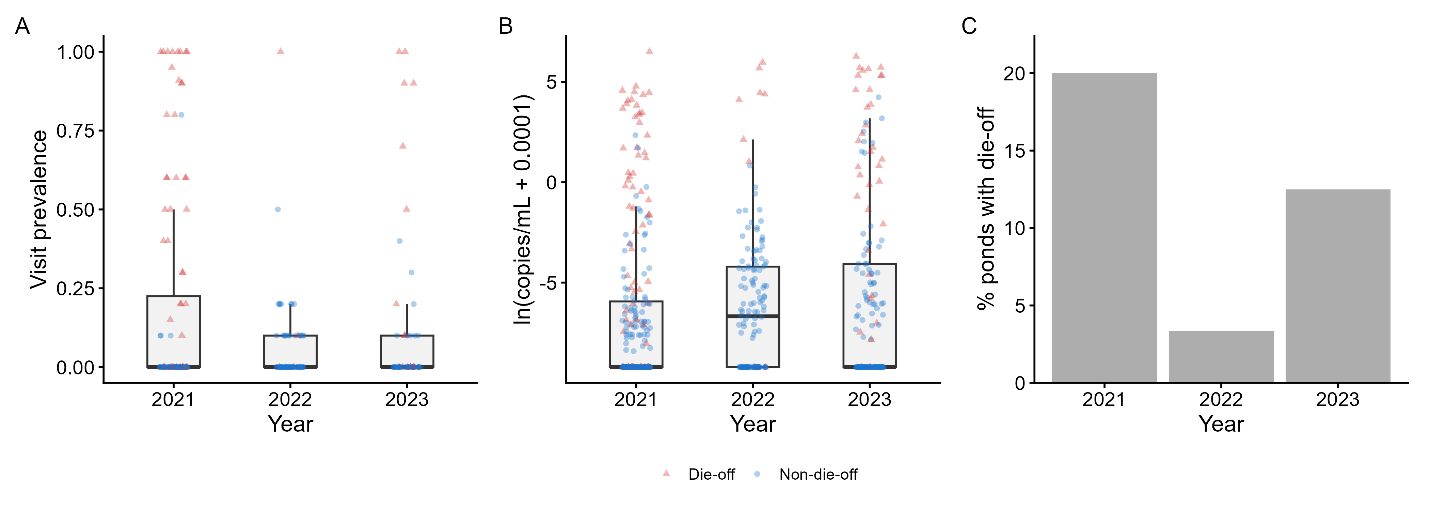

**Figure S2.1.** Inter-annual variation in infection prevalence, environmental viral concentration, and die-off incidence (2021-2023). (A) Infection prevalence at 244 screened visits, including onset but excluding post-onset visits. (B) Environmental viral concentration [ln(copies/mL + 0.0001)] at all 594 visits with observed eDNA, including post-onset visits. Boxplots show medians and interquartile ranges; red triangles indicate die-off pond-years and blue circles indicate non-die-off pond-years. (C) Annual die-off incidence: 8/40 pond-years in 2021 (20.0%), 1/30 in 2022 (3.3%), and 4/32 in 2023 (12.5%).

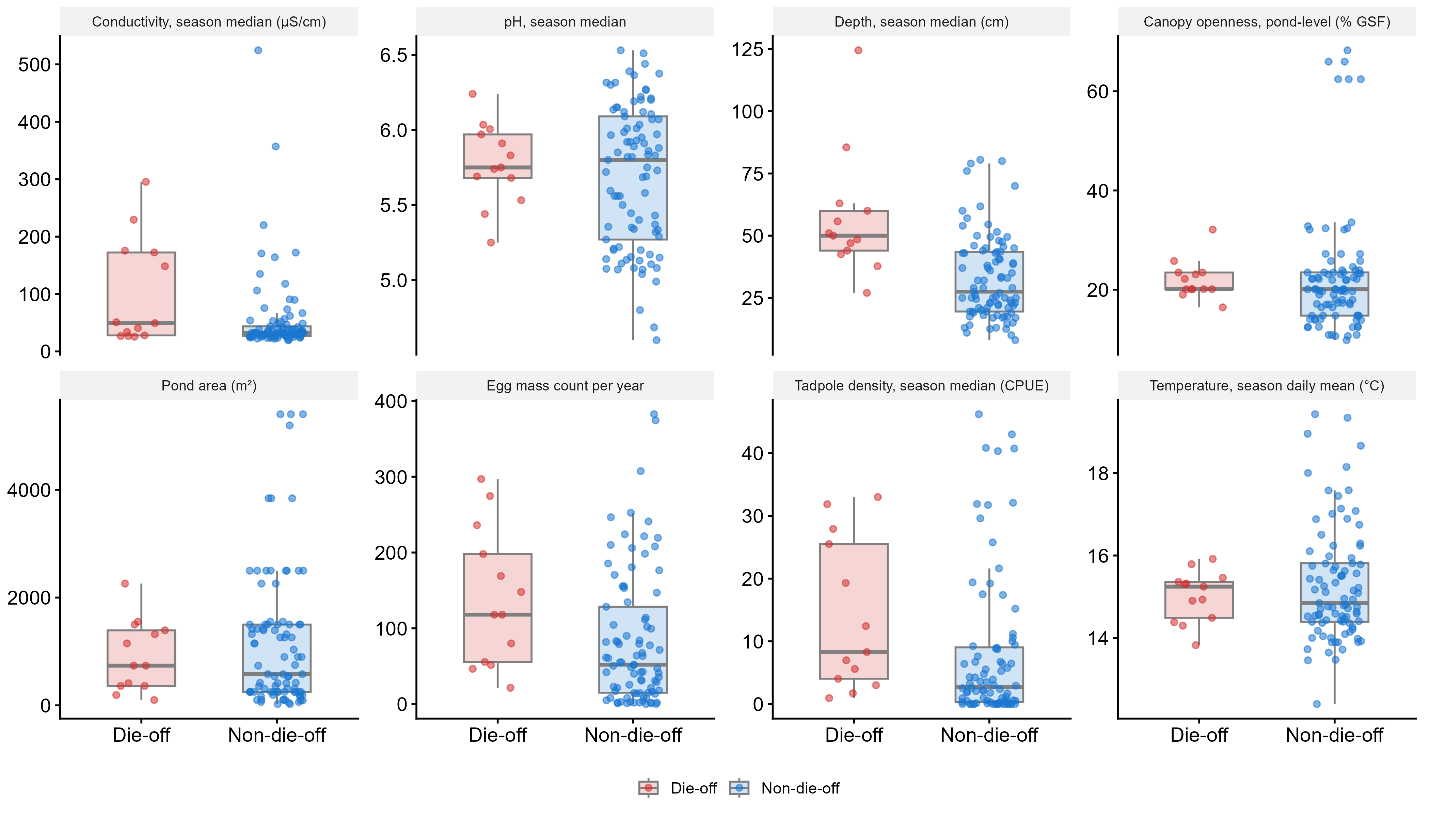

**Figure S2.2.** Environmental and demographic variation between die-off and non-die-off pond-years. Panels show season medians for conductivity, pH, depth, and tadpole density; season mean daily temperature; pond-level canopy openness and area; and annual egg mass count. Tadpole density is expressed as catch per unit effort (CPUE). Points represent pond-years and boxplots show medians and interquartile ranges; red indicates die-off pond-years and blue indicates non-die-off pond-years. The dataset contains 102 pond-years, with density available for 97; canopy and temperature summaries include the imputations described in Appendix S1.

#### S2.1 Ranavirus occurrence

In the parallel eDNA-occurrence screen, conductivity received substantially greater support than the alternatives (Akaike weight = 0.966; ΔAICc ≥ 8.2). Tissue-confirmed ranavirus occurrence (Moran's I = 0.014, p = 0.233), die-off occurrence among ever-infected ponds (I = 0.039, p = 0.228), and residuals from the tissue-occurrence model (I = -0.068, p = 0.787) were not spatially autocorrelated. Distance to the nearest die-off pond was also unsupported as a predictor of tissue-confirmed occurrence (β = 0.33, p = 0.286).

**Table S2.1.** Model comparison for tissue-confirmed ranavirus occurrence. Univariate binomial GLMMs with pond random intercepts were compared on a common sample of 97 pond-years from 38 ponds. The selected conductivity + egg mass count model was refit to all 102 pond-years from 40 ponds (R^2^m = 0.262, R^2^c = 0.369).

| **Model** | **AICc** | **ΔAICc** | **Akaike weight** |
| --- | --- | --- | --- |
| Conductivity | 117.1 | 0.0 | 0.467 |
| Egg mass count | 118.9 | 1.8 | 0.191 |
| Depth | 119.1 | 2.0 | 0.174 |
| Elevation | 121.2 | 4.1 | 0.059 |
| Null | 122.8 | 5.8 | 0.026 |
| Density | 123.3 | 6.3 | 0.020 |
| pH | 124.3 | 7.2 | 0.013 |
| Area | 124.6 | 7.5 | 0.011 |
| Mean temperature | 124.6 | 7.5 | 0.011 |
| Canopy | 124.9 | 7.9 | 0.009 |
| Wetland cover | 124.9 | 7.8 | 0.009 |
| Temp SD | 125.0 | 7.9 | 0.009 |

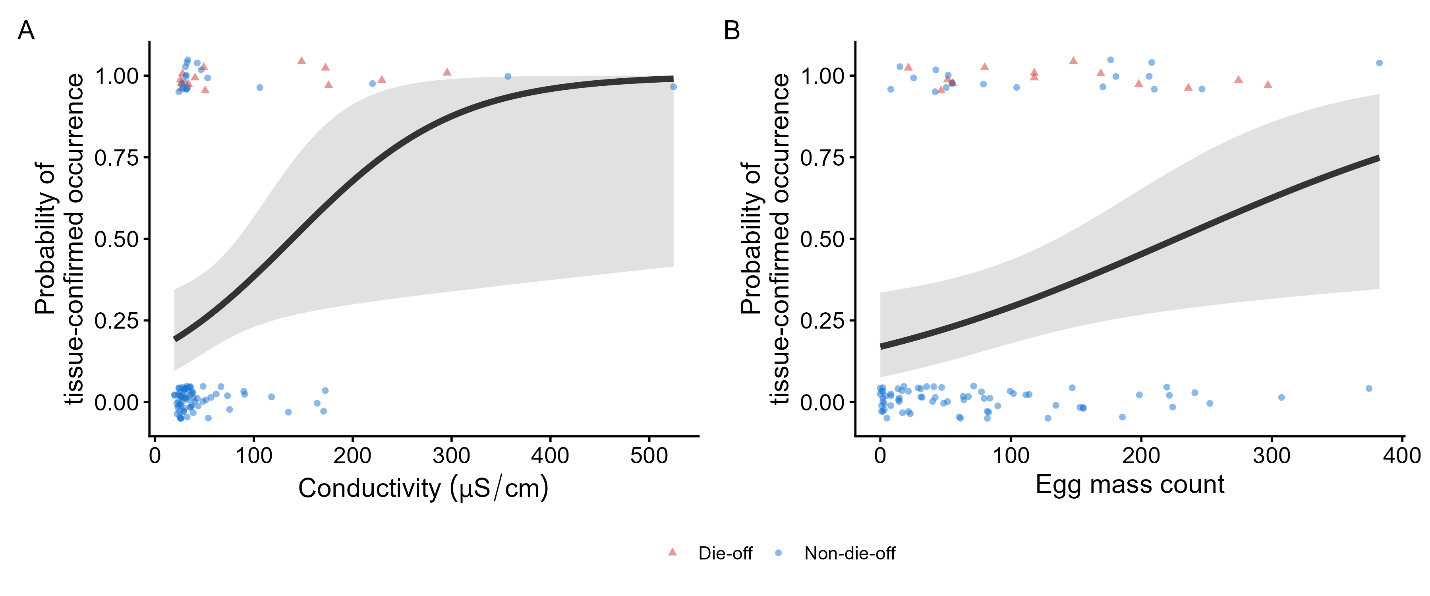

**Figure S2.3.** Marginal effects of conductivity (A) and egg mass count (B) on tissue-confirmed ranavirus occurrence. Curves are predictions from the final binomial GLMM fitted to 102 pond-years from 40 ponds, with the other predictor held at its mean among the modeled pond-years. Lines show fixed-effect predictions with random effects set to zero; ribbons show 95% Wald confidence intervals. Points show jittered binary outcomes; red triangles indicate die-off pond-years and blue circles indicate non-die-off pond-years.

#### S2.2 Infection prevalence

A downsampling sensitivity analysis retained 73 eligible visits in each iteration. Across 500 iterations, the median coefficient for lagged eDNA was 1.85 (empirical 95% range = 1.76-1.98), and the median coefficient for lagged prevalence was 1.25 (1.24-1.38); both predictors remained significant in every iteration. The aggregate LOOCV R^2^ of 0.40 primarily reflects discrimination of infection-positive and infection-free visits, and among infection-positive visits alone, predictive performance was weak (R^2^ = 0.02).

**Table S2.2.** Model comparison for visit-level infection prevalence among ever-infected pond-years. Ten candidate models were fit to a common sample of 95 visits from 25 pond-years using binomial GLMMs with pond-year and observation-level random effects.

| **Model** | **AICc** | **ΔAICc** | **Weight** | ***R^2^_m_*** |
| --- | --- | --- | --- | --- |
| Viral (eDNA_t-1_ + prevalence_t-1_) | 320.7 | 0.0 | 0.667 | 0.504 |
| Viral + Seasonal | 322.1 | 1.4 | 0.331 | 0.567 |
| Viral: prev_t-1_ only | 332.6 | 11.9 | 0.002 | 0.465 |
| Viral: eDNA_t-1_ only | 335.2 | 14.5 | <0.001 | 0.337 |
| Seasonal (DOY) | 343.7 | 23.0 | <0.001 | 0.258 |
| Seasonal + Thermal | 345.6 | 24.8 | <0.001 | 0.261 |
| All environmental | 347.5 | 26.8 | <0.001 | 0.296 |
| Thermal | 358.2 | 37.5 | <0.001 | 0.180 |
| Density | 365.4 | 44.7 | <0.001 | 0.202 |
| Null | 374.9 | 54.2 | <0.001 | 0.000 |

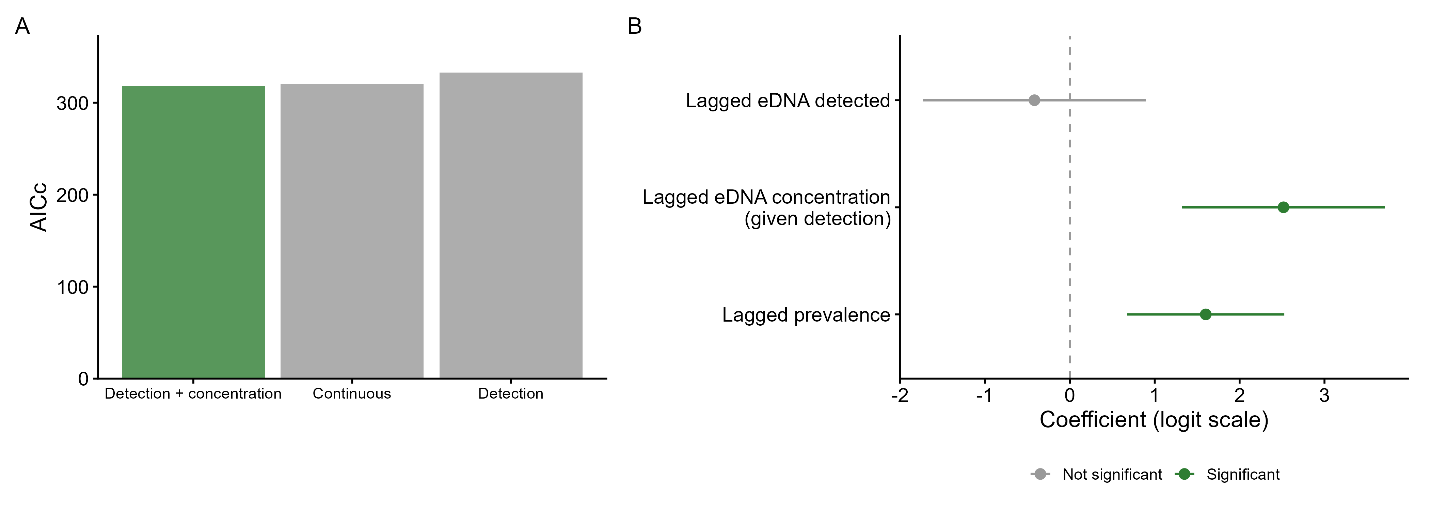

**Figure S2.4.** Alternative parameterizations of lagged eDNA as predictors of subsequent infection prevalence. (A) AICc comparison of models representing eDNA as binary detection, detection plus concentration conditional on detection, or continuous ln(eDNA), fit to the primary common sample (N = 95 visits, 25 pond-years). (B) Coefficients and 95% Wald confidence intervals from the detection-plus-concentration model.

#### S2.3 Viral load

Binary classification using the GMM-derived low-/high-intensity threshold produced consistent results (lagged eDNA: β = 4.17, SE = 1.57, p = 0.008; lagged prevalence: β = 3.13, SE = 1.57, p = 0.046).

**Table S2.3.** Model comparison for viral load [ln(copies/ng DNA + 0.006)] among infected tadpoles. Eleven candidate LMMs were fit to a common sample of 245 individuals from 42 visits and 16 pond-years, with pond-year and visit random intercepts.

| **Model** | **AICc** | **ΔAICc** | **Weight** | ***R^2^_m_*** |
| --- | --- | --- | --- | --- |
| Viral (eDNA_t-1_ + prevalence_t-1_) | 1346.0 | 0.0 | 0.706 | 0.391 |
| Viral + Seasonal | 1348.0 | 2.0 | 0.261 | 0.397 |
| Viral: eDNA_t-1_ only | 1352.8 | 6.8 | 0.024 | 0.320 |
| Viral: prevalence_t-1_ only | 1355.0 | 9.0 | 0.008 | 0.251 |
| Viral: viral load_t-1_ only | 1359.2 | 13.1 | 0.001 | 0.210 |
| Seasonal (DOY) | 1362.1 | 16.1 | <0.001 | 0.167 |
| Seasonal + Thermal | 1364.0 | 18.0 | <0.001 | 0.171 |
| Thermal | 1366.9 | 20.9 | <0.001 | 0.087 |
| All environmental | 1370.3 | 24.3 | <0.001 | 0.174 |
| Null | 1370.4 | 24.3 | <0.001 | 0.000 |
| Density | 1370.9 | 24.9 | <0.001 | 0.031 |

#### S2.4 Environmental viral concentration

In the separate analysis of eDNA detection (N = 130 visits, 31 pond-years), prior detection was strongly associated with subsequent detection (β = 2.39, p < 0.001), whereas lagged infection prevalence was not (β = 0.15, p = 0.660).

Lagging the environmental predictors did not alter the primary conclusions. Lagged pH improved the infection-prevalence model (ΔAICc = -2.2, p = 0.030) and lagged depth improved the environmental viral-concentration model (ΔAICc = -3.7, p = 0.011), but the viral-state predictors remained supported in both models; no lagged environmental predictor improved the viral-load model by >2 AICc units.

**Table S2.4.** Model comparison for environmental viral concentration [ln(copies/mL + 0.0001)] among eDNA-positive visits. Nine candidate LMMs were fit to a common sample of 139 visits from 62 pond-years, with pond-year random intercepts.

| **Model** | **AICc** | **ΔAICc** | **Weight** | ***R^2^_m_*** |
| --- | --- | --- | --- | --- |
| Viral (eDNA_t-1_ + prevalence_t-1_) | 626.0 | 0.0 | 0.721 | 0.630 |
| Viral + Seasonal | 627.9 | 1.9 | 0.279 | 0.631 |
| Autocorrelation ( eDNA_t-1_ only) | 646.5 | 20.5 | <0.001 | 0.564 |
| Shedding (prevalence_t-1_ only) | 704.3 | 78.4 | <0.001 | 0.343 |
| Thermal | 737.2 | 111.2 | <0.001 | 0.151 |
| Seasonal (DOY) | 739.2 | 113.2 | <0.001 | 0.143 |
| All environmental | 740.2 | 114.2 | <0.001 | 0.187 |
| Null | 758.2 | 132.3 | <0.001 | 0.000 |
| Density | 758.4 | 132.4 | <0.001 | 0.017 |

**Table S2.5.** Environmental and host predictors added individually to the core viral-state models for infection prevalence, viral load, and environmental viral concentration. ΔAICc is relative to the corresponding viral-state model refit to the same complete-case subset; p-values refer to the added predictor.

| **Predictor** | **Infection prevalence (prevalence_t_) ΔAICc (p)** | **Viral load (viral load_t_) ΔAICc (p)** | **Environmental viral concentration (eDNA_t_) ΔAICc (p)** |
| --- | --- | --- | --- |
| Stage residuals | -4.6 (0.009) | +1.2 (0.332) | -- |
| Gosner stage | -1.2 (0.033) | +1.4 (0.399) | -- |
| Depth | -0.4 (0.101) | -0.8 (0.087) | -2.1 (0.039) |
| Temperature (7-day mean) | -0.5 (0.044) | +1.7 (0.501) | +1.9 (0.609) |
| Temperature residuals | -- | +1.7 (0.535) | -0.5 (0.098) |
| Temperature SD (7-day) | +1.0 (0.261) | +1.7 (0.535) | +2.1 (0.838) |
| Density | +1.4 (0.361) | +2.0 (0.744) | +1.9 (0.617) |
| Conductivity | +2.2 (0.801) | +1.7 (0.518) | +1.6 (0.429) |
| pH | +2.3 (0.928) | +1.9 (0.644) | +2.0 (0.682) |
| Dissolved oxygen | -- | +1.1 (0.306) | -- |
| DOY | -- | -- | +1.9 (0.587) |

**Table S2.6.** Sensitivity of the three core viral-state models to exclusion of die-off-onset visits. Predictors measured at the previous visit (*t*-1) predict the response at visit *t*.

| **Response** | **N full / no onset** | **Predictor** | **Full β, p** | **No-onset β, p** |
| --- | --- | --- | --- | --- |
| Infection prevalence | 95 / 82 | eDNA*_t−1_* | 1.58, <0.001 | 0.08, 0.881 |
|  |  | Prevalence*_t−1_* | 1.62, <0.001 | 1.85, 0.001 |
| Viral load | 245 / 147 | eDNA*_t−1_* | 2.51, <0.001 | -0.03, 0.955 |
|  |  | Prevalence*_t−1_* | 1.91, 0.002 | 0.22, 0.338 |
| Environmental viral concentration | 139 / 127 | eDNA*_t−1_* | 2.70, <0.001 | 1.98, <0.001 |
|  |  | Prevalence*_t−1_* | 1.36, <0.001 | 1.08, <0.001 |

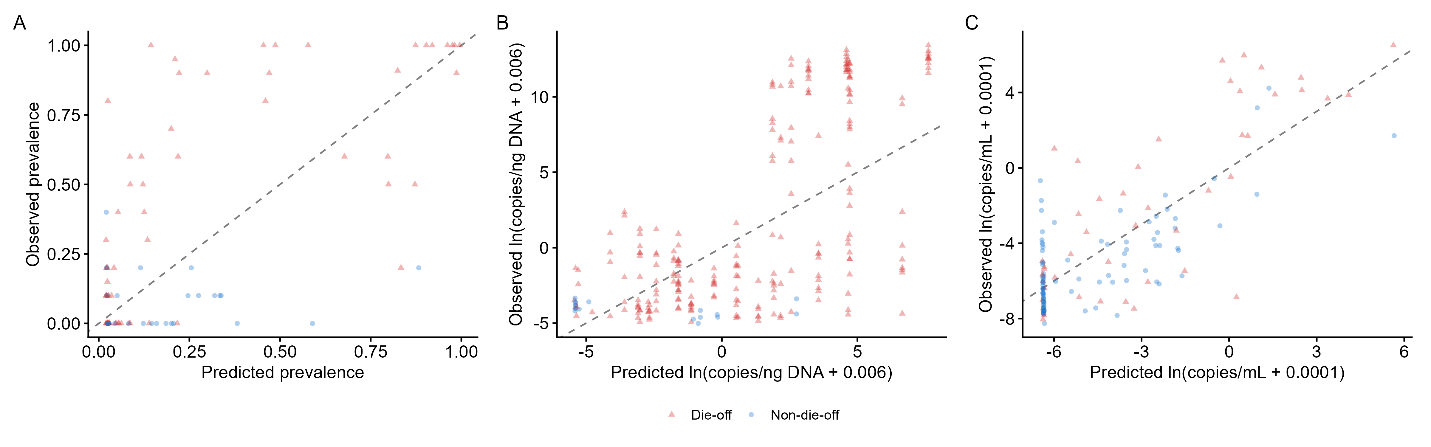

**Figure S2.5.** Leave-one-pond-year-out cross-validation for the three core viral-state models: (A) infection prevalence, (B) viral load among infected tadpoles, and (C) environmental viral concentration among eDNA-positive visits. Points show observed versus fixed-effect predictions for held-out pond-years; the dashed line indicates 1:1 agreement. Panels contain 95 visits, 245 infected individuals, and 139 eDNA-positive visits, and predictive R^2^ values are 0.399, 0.413, and 0.606, respectively.

#### S2.5 Directional time-lagged relationships

After excluding die-off-onset visits, prevalence remained associated with subsequent environmental viral concentration (β = 0.95, p = 0.048), whereas environmental viral concentration did not predict subsequent prevalence or viral load (both p = 0.928). After exclusion of die-off-onset visits (N = 27), global model fit remained adequate (Fisher’s C = 7.65, p = 0.105), but the eDNA→infection prevalence and eDNA →viral load path estimates changed sign.

**Table S2.7.** Directional time-lagged relationships among environmental viral concentration, infection prevalence, and viral load. Each test evaluates whether the predictor at visit *t*-1 predicts the response at visit *t*, accounting for the response at *t*-1 and day of year. P-values were adjusted across the six tests using the Holm method.

| **Direction** | **N** | **β** | **Wald p** | **Holm p** |
| --- | --- | --- | --- | --- |
| eDNA_t-1_ to prevalence_t_ | 97 | 1.57 | <0.001 | <0.001 |
| Prevalence_t-1_ to eDNA_t_ | 91 | 1.21 | <0.001 | 0.001 |
| eDNA_t-1_ to viral load_t_ | 44 | 2.80 | 0.002 | 0.010 |
| Viral load_t-1_ to eDNA_t_ | 60 | 0.98 | 0.109 | 0.109 |
| Prevalence_t-1_ to viral load_t_ | 44 | 2.10 | 0.015 | 0.044 |
| Viral load_t-1_ to prevalence_t_ | 64 | 1.28 | 0.043 | 0.085 |

**Table S2.8.** Path coefficients for the selected piecewise structural equation model (N = 36 visits from 15 pond-years). Predictors were measured at visit *t*−1 and responses at visit *t*. The model included bidirectional eDNA–infection prevalence paths, eDNA_t-1_ → viral load_t_, infection prevalence _t-1_ → viral load _t_, and temporal-persistence terms for all three responses. Global model fit was adequate (Fisher’s C = 3.33, df = 4, p = 0.504).

| **Path** | **Std. beta** | **β** | **p** |
| --- | --- | --- | --- |
| eDNA_t-1_ to prevalence_t_ | 0.20 | 0.65 | 0.114 |
| Prevalence_t-1_ to eDNA_t_ | 0.35 | 1.35 | 0.003 |
| eDNA_t-1_ to viral load_t_ | 0.32 | 2.25 | 0.010 |
| Prevalence_t-1_ to viral load_t_ | 0.38 | 2.24 | 0.003 |
| Prevalence_t-1_ to prevalence_t_ | 0.56 | 1.50 | <0.001 |
| eDNA_t-1_ to eDNA_t_ | 0.60 | 2.76 | <0.001 |
| Viral load_t-1_ to viral load_t_ | 0.30 | 2.65 | 0.021 |

#### S2.6 Die-off discrimination and viral trajectories

The association between first-detection day of year and prevalence slope was concentrated within die-off pond-years (r = −0.61, p = 0.036; N = 12) and was absent among non-die-off pond-years (r = −0.04, p = 0.875; N = 16). Among ever-infected pond-years, lagged eDNA discriminated die-off-onset visits from other visits with an AUC of 0.949 (95% pond-year cluster-bootstrap CI: 0.892-0.990; N = 174 visits, 13 onset events). This analysis distinguishes the timing of onset, not whether a pond-year will ultimately experience a die-off.

**Table S2.9.** Univariate Firth penalized logistic regressions of die-off occurrence against viral trajectory metrics among the 32 ever-infected pond-years (13 die-offs).

| **Predictor** | **Ever-infected: b (p)** | **N** |
| --- | --- | --- |
| Prevalence slope | 2.08 (<0.001) | 28 |
| eDNA slope | 1.51 (0.007) | 17 |
| Peak prevalence | 0.97 (0.003) | 32 |
| Cumulative eDNA | 0.45 (0.219) | 32 |
| eDNA peak | -0.05 (0.750) | 32 |

**Table S2.10.** Univariate linear regressions of prevalence and eDNA trajectory slopes against candidate predictors among ever-infected pond-years. Each row represents a separate model. Prevalence slope was calculated for pond-years with at least three tissue-screened visits and eDNA slope for pond-years with at least three eDNA-positive visits. Predictors were z-standardized before analysis.

| **Predictor** | **Prevalence slope: β (p)** | **N** | **eDNA slope: β (p)** | **N** |
| --- | --- | --- | --- | --- |
| *Static* | | | | |
| Pond area | −0.006 (0.075) | 28 | −0.045 (0.058) | 17 |
| Median pH | 0.004 (0.072) | 28 | 0.003 (0.928) | 17 |
| Egg mass count | 0.003 (0.101) | 28 | −0.030 (0.290) | 17 |
| Median depth | 0.002 (0.163) | 28 | 0.009 (0.718) | 17 |
| Canopy cover | 0.006 (0.161) | 28 | −0.018 (0.490) | 17 |
| Elevation | −0.002 (0.399) | 28 | −0.028 (0.341) | 17 |
| Conductivity | 0.000 (0.800) | 28 | −0.000 (0.970) | 17 |
| *First detection* | | | | |
| First detection DOY | −0.004 (0.011) | 28 | −0.018 (0.651) | 17 |
| First detection temperature | −0.003 (0.101) | 28 | −0.005 (0.861) | 17 |
| First detection density | 0.002 (0.172) | 28 | 0.086 (0.029) | 17 |
| First detection eDNA | 0.003 (0.178) | 28 | −0.004 (0.864) | 17 |
| First detection prevalence | −0.001 (0.483) | 26 | −0.038 (0.069) | 16 |
| *Seasonal* | | | | |
| Season temperature SD | 0.004 (0.106) | 28 | −0.013 (0.638) | 17 |
| Season mean temperature | −0.001 (0.705) | 28 | −0.029 (0.236) | 17 |

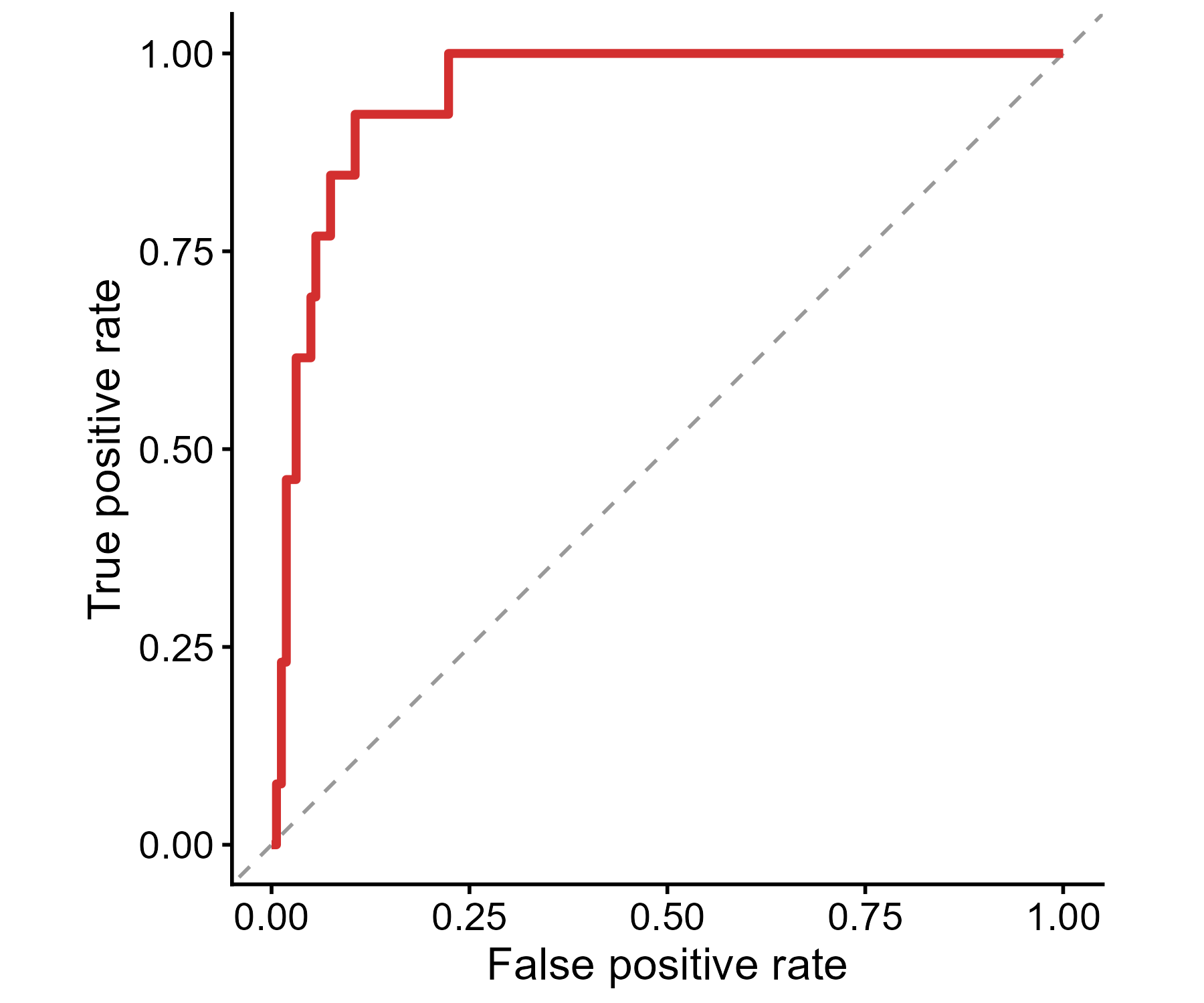

**Figure S2.6.** ROC curve for discrimination of die-off-onset visits from other visits using lagged eDNA among ever-infected pond-years (N = 174 visits, 13 onset events). AUC = 0.949 (95% pond-year cluster-bootstrap CI: 0.892-0.990).

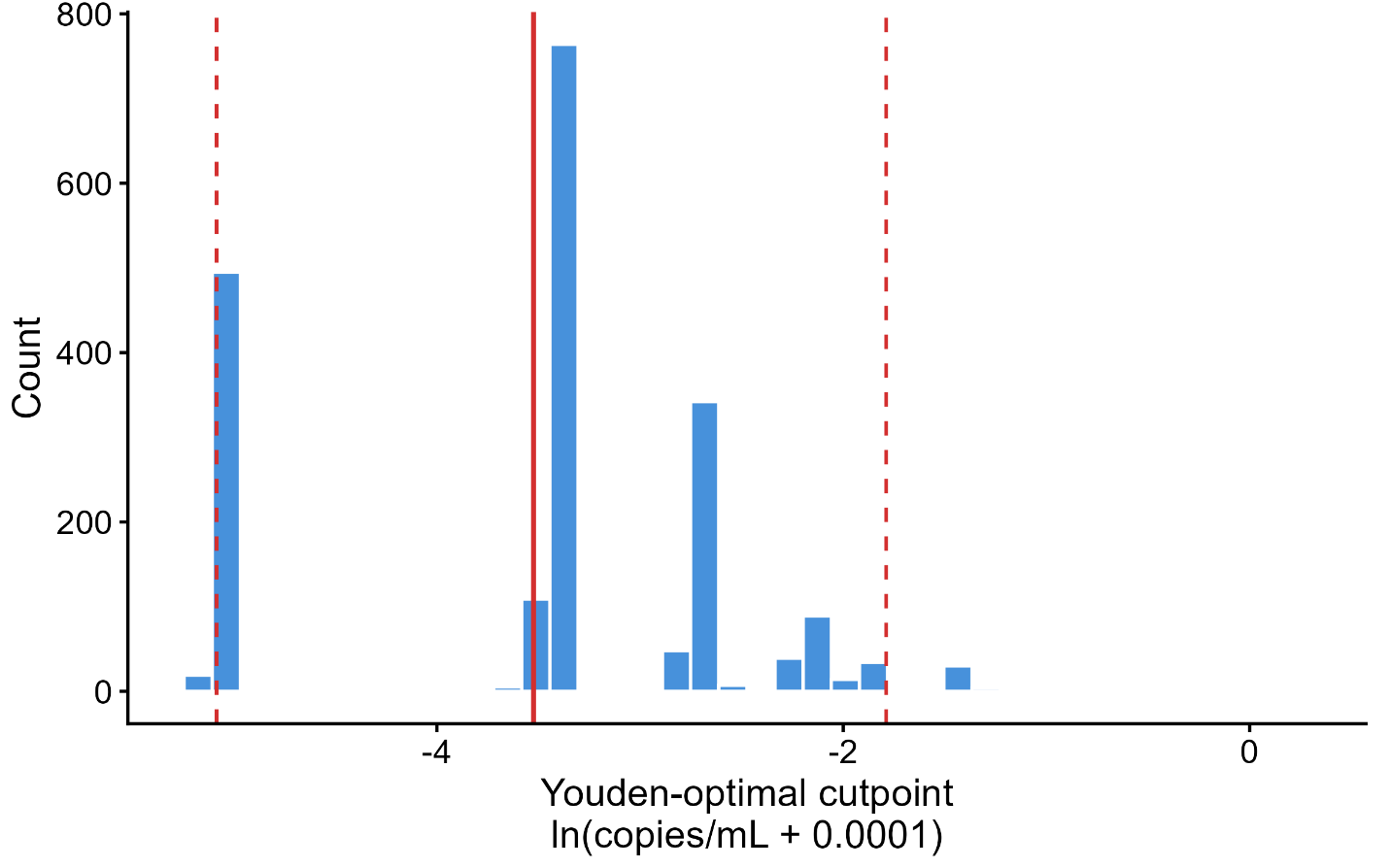

**Figure S2.7**. Pond-year cluster-bootstrap distribution of the Youden-optimal lagged-eDNA cutpoint for discriminating die-off-onset visits among ever-infected pond-years (2,000 resamples). The estimated cutpoint varied substantially among bootstrap samples. Cutpoints are expressed as ln(copies/mL + 0.0001). The solid red line marks the bootstrap mean (-3.52); dashed red lines mark the 95% percentile interval (-5.08 to -1.79).

#### S2.7 Hall et al. (2018) cross-dataset comparison

Data from the Hall et al. (2018) dataset comprised 8 ponds (4 die-off and 4 non-die-off) sampled during a single 2014 season at Yale-Myers Forest.

For infection prevalence (N = 27 visits, 8 ponds), seasonal and temperature predictors received greater support than viral-state predictors alone (ΔAICc = 8.3), although adding lagged eDNA to the seasonal and temperature model produced a competitive model (ΔAICc = 0.5).

For viral load (N = 25 visits), viral-state predictors received substantially greater support than seasonal and temperature predictors (ΔAICc = 13.9; Table S2.11), consistent with the primary dataset.

**Table S2.11.** Model comparison for mean viral load in the Hall et al. (2018) dataset. Thirteen candidate LMMs were fit to 25 visits from eight ponds with a pond random intercept.

| **Model** | **AICc** | **ΔAICc** | **Weight** |
| --- | --- | --- | --- |
| Viral + Seasonal | 66.0 | 0.0 | 0.269 |
| Viral + Thermal | 66.1 | 0.1 | 0.261 |
| Viral: eDNA_t-1_ + prevalence_t-1_ | 66.2 | 0.2 | 0.242 |
| Viral + Seasonal + Thermal | 67.6 | 1.7 | 0.118 |
| eDNA_t-1_ + Seasonal + Thermal | 68.5 | 2.5 | 0.077 |
| eDNA_t-1_ + Seasonal | 70.6 | 4.6 | 0.027 |
| Viral: eDNA_t-1_ | 74.2 | 8.2 | 0.004 |
| Viral: viral load_t-1_ | 77.2 | 11.2 | 0.001 |
| Seasonal + Thermal | 80.1 | 14.1 | 0.000 |
| Seasonal | 80.9 | 14.9 | 0.000 |
| Viral: prevalence_t-1_ | 81.8 | 15.8 | 0.000 |
| Thermal | 86.9 | 21.0 | 0.000 |
| Null | 93.0 | 27.0 | 0.000 |

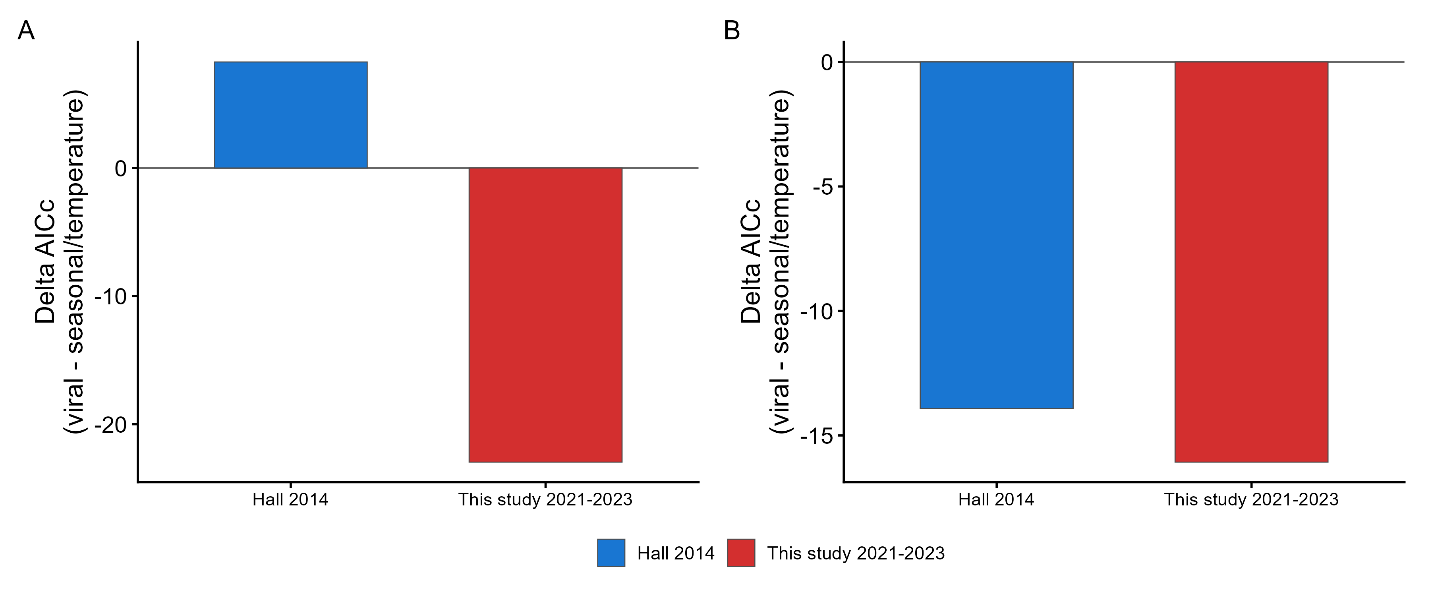

**Figure S2.8.** Cross-dataset comparison of support for viral-state versus seasonal and temperature predictors in the current study (2021-2023) and the 2014 Hall et al. (2018) dataset, both from Yale-Myers Forest. Panels show infection prevalence (A) and viral load (B). Bars show AICc for the best viral-state model minus AICc for the best seasonal/temperature model within each dataset and response; negative values favor viral-state predictors. Viral-state predictors received greater support for viral load in both datasets, whereas support for prevalence differed between datasets.

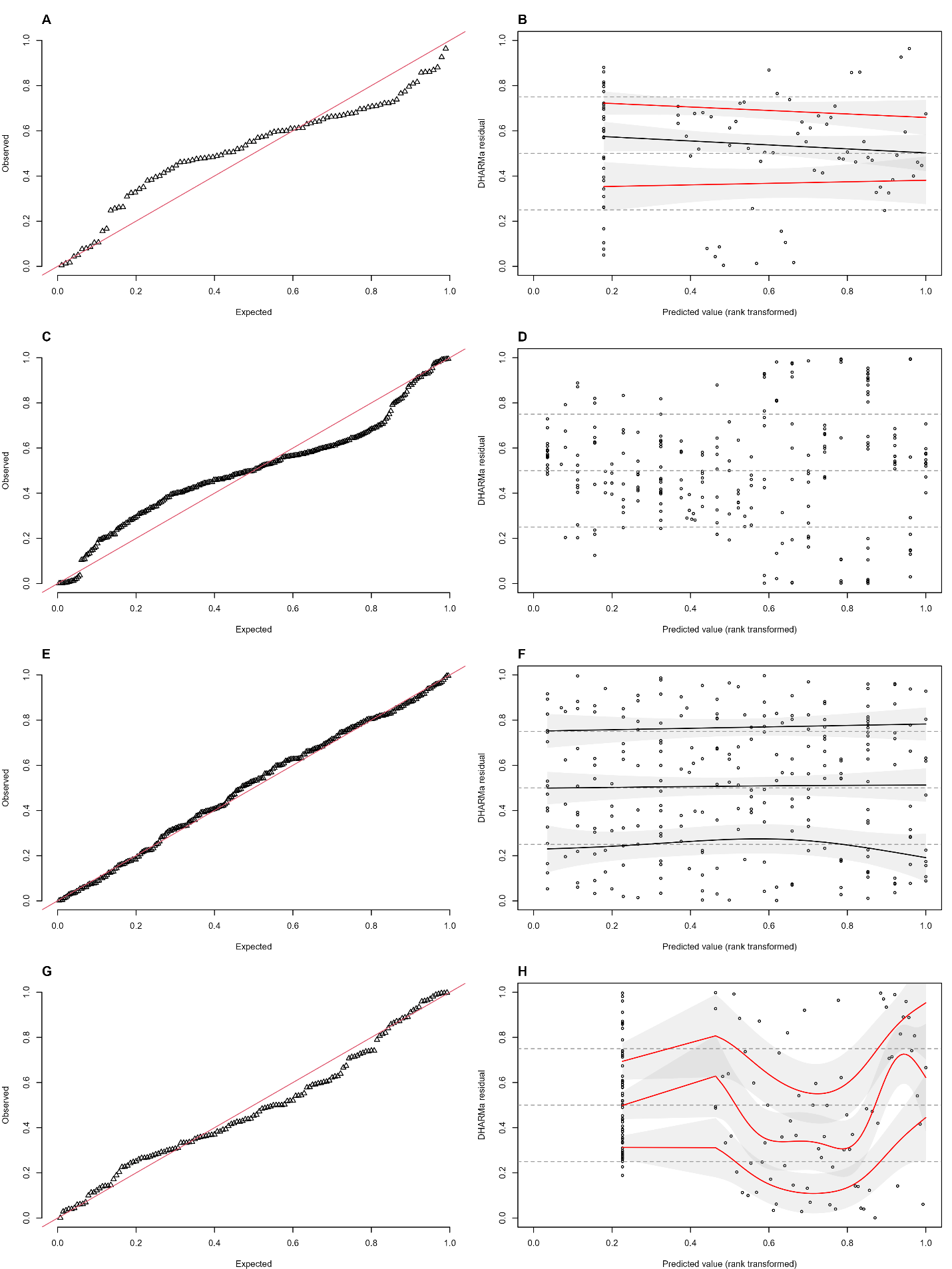

**Figure S2.9.** DHARMa scaled residual diagnostics for infection prevalence (A–B), viral load among infected tadpoles (C–D), binary low-/high-intensity infection classification (E–F), and environmental viral concentration (G–H). Each row shows a QQ plot (left) and residuals versus rank-transformed fitted values (right), based on 1,000 conditional simulations. Dashed horizontal lines mark expected residual quartiles; solid curves show quantile fits. In the residual-versus-fitted panels, gray bands show approximate 95% intervals, and red quantile curves indicate departures after BH adjustment. Viral-load quantile curves are omitted because those fits produced convergence warnings.
